# Color-dependent foraging in *C. elegans* integrates chromoprotein photosensitization with bacterial metabolic cues

**DOI:** 10.64898/2026.08.27.747292

**Authors:** Rohil Hameed, Vedat Sari, Yang Yue, Zhenwen Yu, Sergei Koshkin, Charles Evans, Andrey Parkhitko, Scott F. Leiser, Alaattin Kaya

**Affiliations:** ¹School of Medicine, Department of Biochemistry and Molecular Genetics, University of Virginia, Charlottesville, VA, USA; School of Medicine, Department of Biochemistry and Molecular Biology, Virginia Commonwealth University, Richmond, VA; Aging Institute of UPMC and the University of Pittsburgh, Pittsburgh, PA, USA; Department of Human Genetics, School of Public Health, University of Pittsburgh, Pittsburgh, PA, USA; Huck Institutes of the Life Sciences, The Pennsylvania State University, University Park, PA, USA; School of Medicine, Department of Internal Medicine, University of Michigan, Ann Arbor, Michigan, USA; Division of Endocrinology and Metabolism, Department of Medicine, University of Pittsburgh, Pittsburgh, PA, USA; School of Medicine, Department of Molecular and Integrative Physiology, University of Michigan, Ann Arbor, United States

**Keywords:** *C. elegans*, foraging, gut–brain axis, serotonin, neuropeptide signaling

## Abstract

Animals rely on color to navigate complex environments, yet how eyeless organisms use chromatic information to guide food choice remains poorly understood. Here, we show that *Caenorhabditis elegans* exhibits robust color-dependent foraging driven by microbial chromophores, preferentially consuming red while avoiding blue chromoprotein-expressing bacteria across bacterial backgrounds and wild isolates. This discrimination persists in darkness and independently of photoreceptor, revealing a mechanism beyond canonical photoreception. Purified chromoproteins and bacterial metabolite fractions independently reproduce preference, demonstrating complementary chromatic and post-ingestive metabolic cues. Mechanistically, blue chromoproteins generate singlet oxygen, producing oxidative stress and remodeling bacterial tryptophan and pterin metabolism, whereas red food promotes serotonin production and feeding-associated neuropeptide signaling. Disrupting serotonin biosynthesis or neuropeptide processing abolishes color preference. Together, our findings reveal a previously unrecognized, novel sensory strategy in which wavelength-selective pigment photochemistry transforms microbial color into metabolic information that is integrated through gut–brain neuroendocrine signaling to guide foraging behavior in an eyeless animal.

## INTRODUCTION

Color is one of the most informative environmental features guiding critical behaviors such as foraging, survival, and mating in diverse animal species (*1*, *2*). However, the mechanisms by which eyeless organisms extract and utilize spectral information remain poorly defined (*3*, *4*) The nematode *Caenorhabditis elegans* provides a powerful genetic model to dissect these mechanisms. Despite lacking eyes and classical opsins, worms evaluate environmental inputs, including microbial pathogenicity and coloration, to guide foraging decisions (*5*, *6*). For instance, under white light, *C. elegans* avoids harmful, blue-pigmented bacteria by evaluating the spectral ratio of blue to amber wavelengths (*7*, *8*). This opsin-independent color sense complements the function of LITE-1, a noncanonical, gustatory-receptor-like photoreceptor that directly absorbs UV/blue light to mediate photophobic and feeding-inhibition responses(*9*, *10*). Because visible light can inflict physiological stress and reduce lifespan via photo-oxidative damage, avoiding short-wavelength spectral signatures has clear ecological utility (*11*).

Natural microbial communities are visually heterogeneous. Bacteria synthesize pigments spanning the visible spectrum, including carotenoids (yellow-orange-red), phenazines/pyocyanin (blue-green), violacein (purple), and melanins (brown-black), all of which serve essential roles in photosynthesis, photoprotection, redox homeostasis, and microbial antagonism (*12*, *13*) . These pigments alter both the spectral signature reflected to a foraging animal and the chemical milieu encountered upon ingestion (*5*).

Beyond initial sensory detection, behavioral decisions in *C. elegans* are dynamically shaped by neuromodulatory circuits that regulate feeding, locomotion, and foraging states. Monoamines such as serotonin, together with diverse neuropeptide signaling systems, coordinate behavioral transitions in response to food availability and environmental context (*14*). Serotonergic neurons respond rapidly to food-associated cues to promote feeding behaviors, whereas neuropeptide networks modulate sustained locomotor and behavioral states (*15*). Moreover, ingested microbial signals can influence these circuits through intestinal metabolism, providing a mechanism by which the physiological consequences of a food source can modify subsequent behavioral decisions (*16*). However, whether microbial color can influence such neuromodulatory pathways independently of pathogenicity, and how chromatic information might be converted into biochemical signals accessible to an animal lacking conventional visual machinery, remain unknown.

We therefore considered a model in which color-dependent food choice emerges from the integration of chromophore photochemistry with internal metabolic sensing rather than from conventional photoreception alone. In this framework, microbial chromophores could function as wavelength-selective chemical interfaces that convert incident light into distinct local biochemical states. Chromophore-dependent photochemistry could thereby generate extracellular signals, including reactive oxygen species, that provide rapid pre-ingestive information about a potential food source. In parallel, the same photochemical properties could remodel bacterial metabolism, producing distinct metabolite signatures that are encountered after ingestion and communicated from the intestine to the nervous system. Such complementary pre- and post-ingestive mechanisms would allow *C. elegans* to couple the spectral properties of microbial food with its physiological consequences and translate microbial coloration into adaptive foraging decisions.

To directly test this possibility while decoupling pigmentation from bacterial virulence, we engineered non-pathogenic *Escherichia coli* food sources to express non-toxic, coral-derived chromoproteins spanning distinct visible wavelengths. Using this system, we demonstrate that *C. elegans* robustly prefers red chromoprotein-expressing bacteria while avoiding blue, with discrimination persisting across bacterial genetic backgrounds, diverse wild isolates, darkness, and loss of the photoreceptor LITE-1. Biochemical fractionation and purified chromoprotein assays further identify both the chromoprotein itself and chromoprotein-induced bacterial metabolites as instructive cues. Cross-kingdom metabolomic and transcriptomic analyses reveal that blue and red chromoproteins generate divergent oxidative, metabolic, and neuroendocrine states, while genetic and biochemical analyses identify serotonin and neuropeptide signaling as essential components of color-dependent food choice. Together, our findings reveal a previously unrecognized sensory strategy in which microbial chromoprotein photochemistry transforms chromatic information into biochemical signals that are integrated with post-ingestive metabolic cues through gut–brain neuroendocrine pathways to guide foraging behavior in an eyeless animal.

## RESULTS

### Chromophores Derived from Coral Reefs Modulate *C. elegans* Color-dependent Foraging Behavior

*Caenorhabditis elegans* thrives in decomposing organic matter populated by metabolically diverse bacteria (*17*), many producing pigments that create a visually heterogeneous landscape for this eyeless nematode (*18*) . While indirect cues such as bacterial pathogenesis modulate nematode foraging, whether *C. elegans* exhibits intrinsic preferences based solely on visible color remains undetermined.

We therefore engineered *E. coli* (*BL21*), a standard non-pathogenic laboratory food source, to express synthetic or natural coral-derived chromoproteins (**Fig. 1A**) (*19*, *20*), producing distinct blue, red, yellow, orange, purple, and green colors under visible light (**Fig. 1B**). N2 (wild-type) worms reared from embryo to young adult on each strain showed no developmental delay or morphological defect (**Fig. 1C**), but yellow and blue bacteria significantly decreased thrashing rate (*p = 0.001*, student’s t-test) (**Fig. 1D**) and red bacteria increased pharyngeal pumping rate (*p = 0.01*, student’s t-test) relative to the non-chromogenic control (**Fig. 1E**).

**Figure 1.**
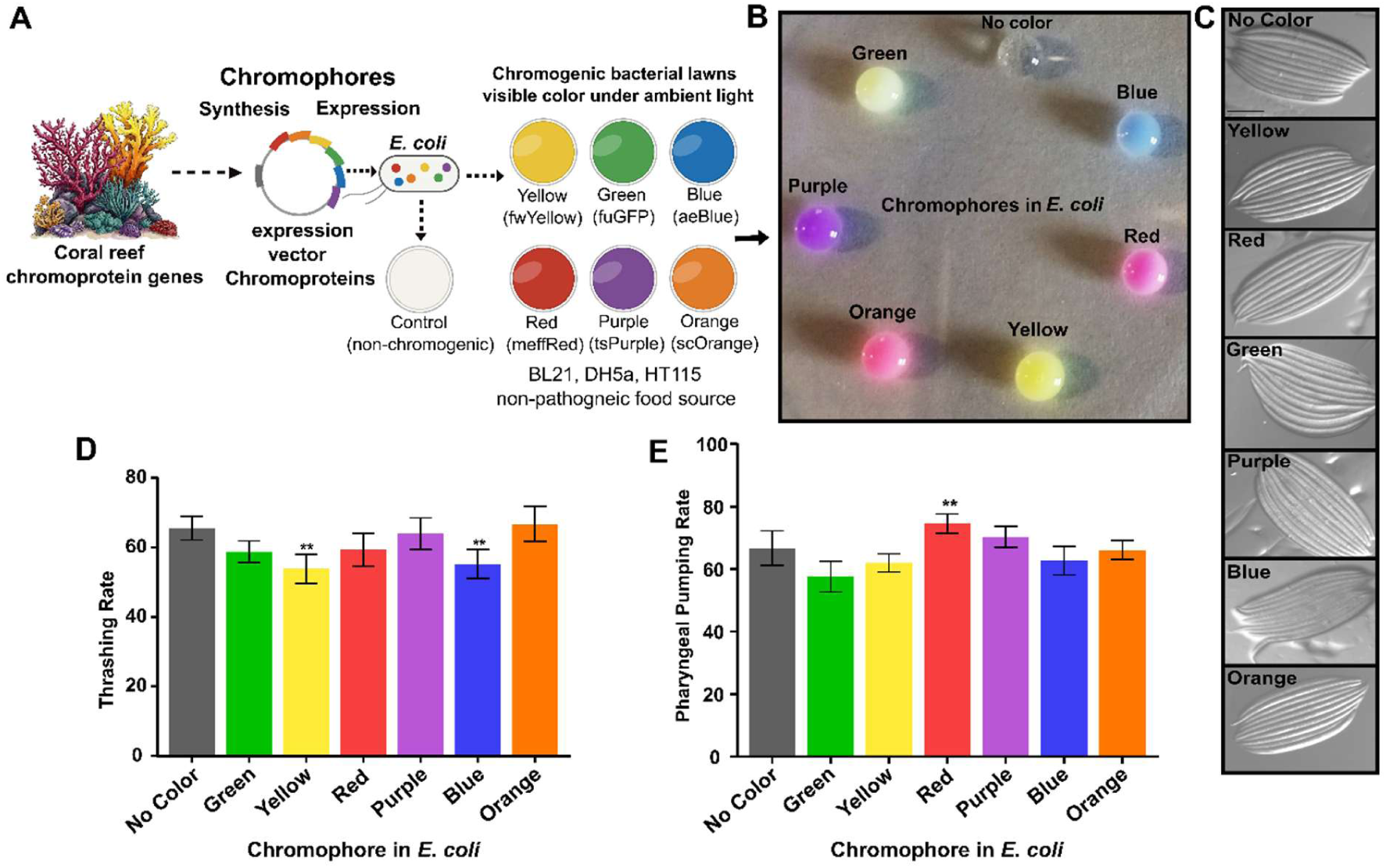
Chromophore expression and underlining behavioral effects on *C. elegans*: **(A)** Schematic depicting the origin of different chromophores from coral reefs and the experimental method for their synthesis in bacteria. **(B)** Representative images show six bacterial cultures of *E. coli (BL21)* expressing different chromophores that the color can be observed in visible light**. (C)** Bright field images of young adult worms (wild-type-N2) fed with *E. coli (BL21)* expressing individual chromophore (n = 10). **(D)** Bar graph shows the thrashing rate of young adult N2 worms after being reared on chromophore bacteria from embryo to young adult stage (n = 20 worms on each plate, \**p = 0.001*). **(E)** Graphs show pharyngeal pumping rate of young adult N2 worms after reared on chromophore bacteria (n = 20, \**p = 0.010*). The data represent three biological replicates for each experiment.

In a choice assay presenting young adult N2 worms simultaneously with all live chromogenic lawns (**Fig. 2A**), ∼30% of animals accumulated on red bacteria within 5 hours under visible light, followed by the non-chromogenic control (∼20%), with the remaining ∼50% distributed across purple, orange, yellow, green, and blue lawns (**Fig. 2B, C**). The bias persisted at 24 and 48 hours, consumption of red lawns was greater than that of the other lawns (**Fig. 2B and Fig. S1A, B**).

**Figure 2.**
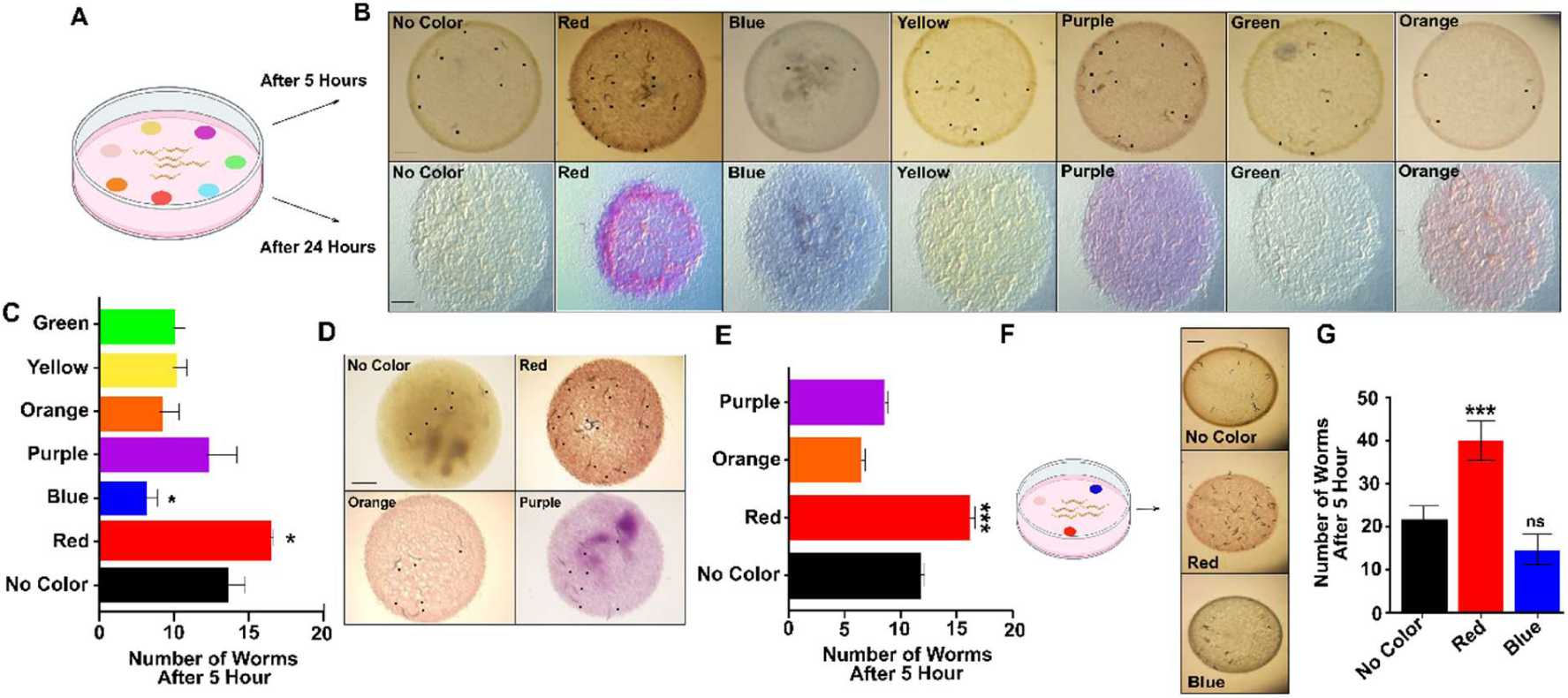
Chromophore-guided foraging behavior in *C. elegans:* **(A)** Schematic depicting the chromophore (bacteria) choice assay arrangement for the young adult N2 worms with different time points. **(B)** Representative stereo microscope images show the chromophore-based foraging behavior of N2 worms after 5-and 24 hours incubation under ambient light conditions. The worms on each bacterial lawn were depicted with a black dot mark. **(C)** Bar graph represents the number of N2 worms on each chromogenic bacterial lawn after 5 hours of incubation (n=100 worms on each plate were assayed, \**p = 0.0133 (red), 0.433 (blue), 0.0240 (yellow), 0.0433 (green)*. **(D)** Representative plate images of **c**hromophore choice of attraction in between red, orange and purple color. The worms on each bacterial lawn were depicted with a black dot mark. **(E)** Bar graph shows the number of worms on red, orange and purple bacterial lawns (n = 100 worms on each plate, \**p = 0.007* (*red*), *0.0121* (orange*), 0.0021 (purple)*. **(F)** Representative plate images show the chromophore attraction of N2 worms upon exposure to no color (control) bacteria, red and blue chromophore bacteria. **(G)** The bar graph represents the number of worms observed on each bacterial lawn after 5 h (\**p* = 0.0068). All assays presented in this figure were performed on standard NGM plates seeded with OP50 bacteria. Data represent three biological replicates for each experiment.

To test whether the preference is specific to red, we assayed red, orange, and purple lawns only, spanning decreasing wavelength ranges (∼620-750, ∼590-620, and ∼380-450 nm, respectively) (*21*). Worms again significantly (*p = 0.0005*, ANOVA test) accumulated on red after 5 hours (**Fig. 2D, E**), and a three-choice assay of colorless, red, and blue lawns confirmed significant preference for red and lowest preference for blue (*p = 0.00068*) (**Fig. 2F**).

The behavior was not strictly light dependent: over 24 h, worms preferentially consumed red bacteria under both ambient-light and dark conditions (**Fig. S2**). Repeating the assay under both conditions with wild isolates from distinct niches, CX11262 (U.S.A.), GXW1 (China), QX2263 (Argentina), and ED3049 (South Africa) (*22*), all significantly accumulated on and preferentially consumed red lawns (*p ≤ 0.05*), whereas blue avoidance was mixed. The CX11262 avoided blue in both light and dark, QX2263 only in light, and GXW1 and ED3049 in neither (**Fig. S3-6**). Red attraction is therefore commonly evolved, while blue avoidance appeared strain-and niche-specific.

We next tested the requirement for LITE-1, which mediates UV light-dependent avoidance of pathogenic bacteria (*23*) . LITE-1 mutants (TQ8245) nonetheless retained significant (*p ≤ 0.05*) red preference and blue avoidance at 5 and 24 hours (**Fig. S7A-B**), indicating that color-biased foraging is LITE-1 independent. The behavior was likewise independent of bacterial background. Worms, cultured on bacterial lawns of red and blue chromophores expressed in *DH5α*, *BL21*, and *HT115* (**Fig. S8**) and each elicited significant red preference (*p = 0.005,* student’s t-test), blue avoidance, and preferential red consumption over 24 and 48 hours (**Fig. 3A, B and Fig. S9**). Finally, a red chromophore of independent origin (meffRed) reproduced this significant attraction (*p = 0.0049*, student’s t-test) (**Fig. S10**).

**Figure 3.**
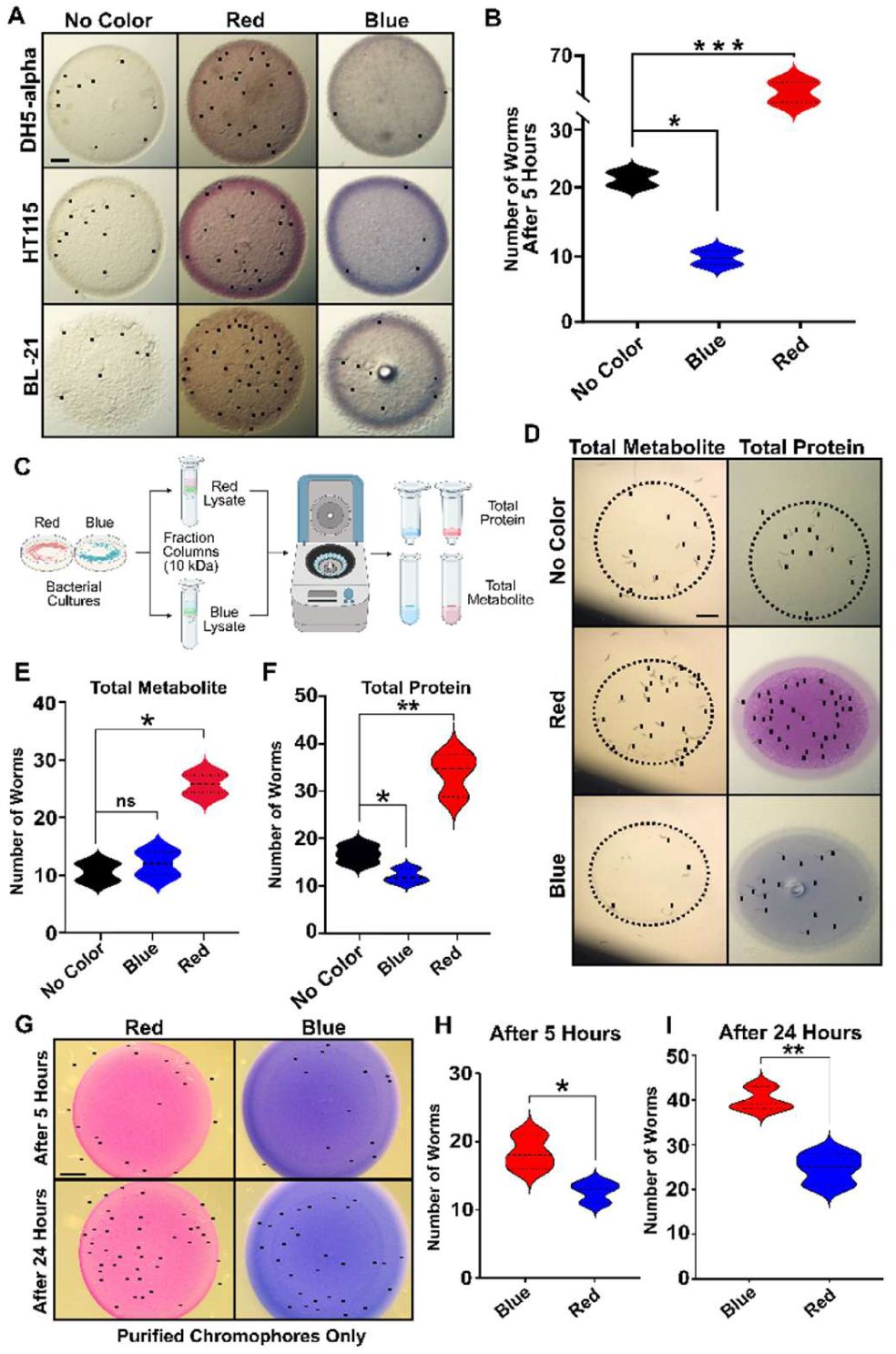
Red chromophore attracts *C. elegans*: **(A)** Plate images represent the chromogenic bacterial lawns of blue and red chromophores, expressed in three different bacterial genetic backgrounds. The worms on each bacterial lawn were depicted with a black dot mark. (**B**) The graph represents the cumulative number of worms from each chromogenic bacteria of different genetic background for no color (control), blue and red attraction after 5 hours of incubation. (\**p = 0.005,* student’s t-test) (**C**) Depiction of column based experimental procedure for fractioning the total bacterial metabolite and protein mixes from red and blue chromogenic bacterial cells. (**D**) Representative plate images of choice assay, seeded with total metabolite or protein mixes isolated from red and blue chromogenic bacterial cells. The worms on each bacterial lawn were depicted with a black dot mark. **(E)** Bar graph represents the number of N2 worms on each metabolite and (**F**) protein mixes. Images were obtained after 5 hours of incubation under ambient light condition. (n = 100 worms on each plate, (\**p = 0.0014, 0.0329,* student’s t-test). **(G)** Images of plates, seeded with purified red and blue chromophores used for choice assay at two different time point (5 and 24-hour). The worms on each lawn were depicted with a black dot mark. (**H**) The graphs show the number of N2 worms after 5 hours of incubation on plates seeded with purified chromophores of red and blue proteins. (n = 100 worms on each plate were assayed, (\**p = 0.005,0.0048,* student’s t-test). All assays presented in this figure were done on standard NGM plates with HT115 bacteria. The data represent three biological replicates for each experiment.

Collectively, these results provide unbiased evidence that an eyeless nematode discriminates among colors, favoring red and avoiding blue, and that color cues guide foraging decisions independent of bacterial pathogenicity and of a canonical visual system.

### *C. elegans* integrates both color and color-associated bacterial metabolite cues to guide foraging decisions

We next asked whether this behavior is driven by color itself or by chromoprotein-induced bacterial metabolites that initiate chemotaxis. Red and blue chromophore-expressing *HT115* lysates were fractionated on a 10-kDa column into metabolite and protein mixes (**Fig. 3C**); both chromophores exceed 10 kDa and were retained in the 1X-PBS-washed protein fraction. Equal amounts of each were seeded onto NGM plates (**Fig. 3D**). N2 worms significantly preferred both the red protein fraction (*p = 0.0014*, student’s t-test) and the red metabolite fraction (*p = 0.0329*, student’s t-test) over their blue counterparts (**Fig. 3E, F**). Identical results were obtained with fractions from *OP50* and *BL21* and in the LITE-1 mutant (**Fig. S11-13**), indicating that both color and chromophore-induced metabolic changes mediate foraging, independent of bacterial origin and LITE-1.

Because these protein fractions are heterogeneous, we purified His-tagged red and blue chromophores and seeded the pure proteins onto NGM plates (**Fig. 3G, H**). Strikingly, worms showed significant preference for purified red over blue chromophore at both 5 hours (*p = 0.005*) and 24 hours (*p = 0.0048*, student’s t-test) (**Fig. 3G-I**). Visible color is therefore sufficient to drive foraging preference in eyeless *C. elegans*, with chromatic and metabolic signals acting as dominant, instructive cues.

### Cross-species multi-Omics reveal a molecular basis for color-dependent food discrimination in *C. elegans*

To gain molecular insight, we combined bacterial metabolomics with host transcriptomics. Untargeted metabolomics of *HT115* expressing red or blue chromoproteins alongside non-chromogenic controls detected 1,134 mass features, 611 putatively annotated (**File S1**). PCA of log₂-transformed, Pareto-scaled intensities separated all three groups, PC1 (58.5%) separating chromoprotein-expressing bacteria from control and PC2 (18.3%) distinguishing red from blue (**Fig. 4A**). Blue bacteria showed the strongest displacement along PC1 and a larger differential response (109 metabolites up, 136 down) than red (71 up, 67 down), with 68 up and 95 down unique to blue versus 30 and 26 unique to red (41 shared in each direction) (**Fig. 4B**), indicating that aeBlue imposes the broader metabolic perturbation.

**Figure 4.**
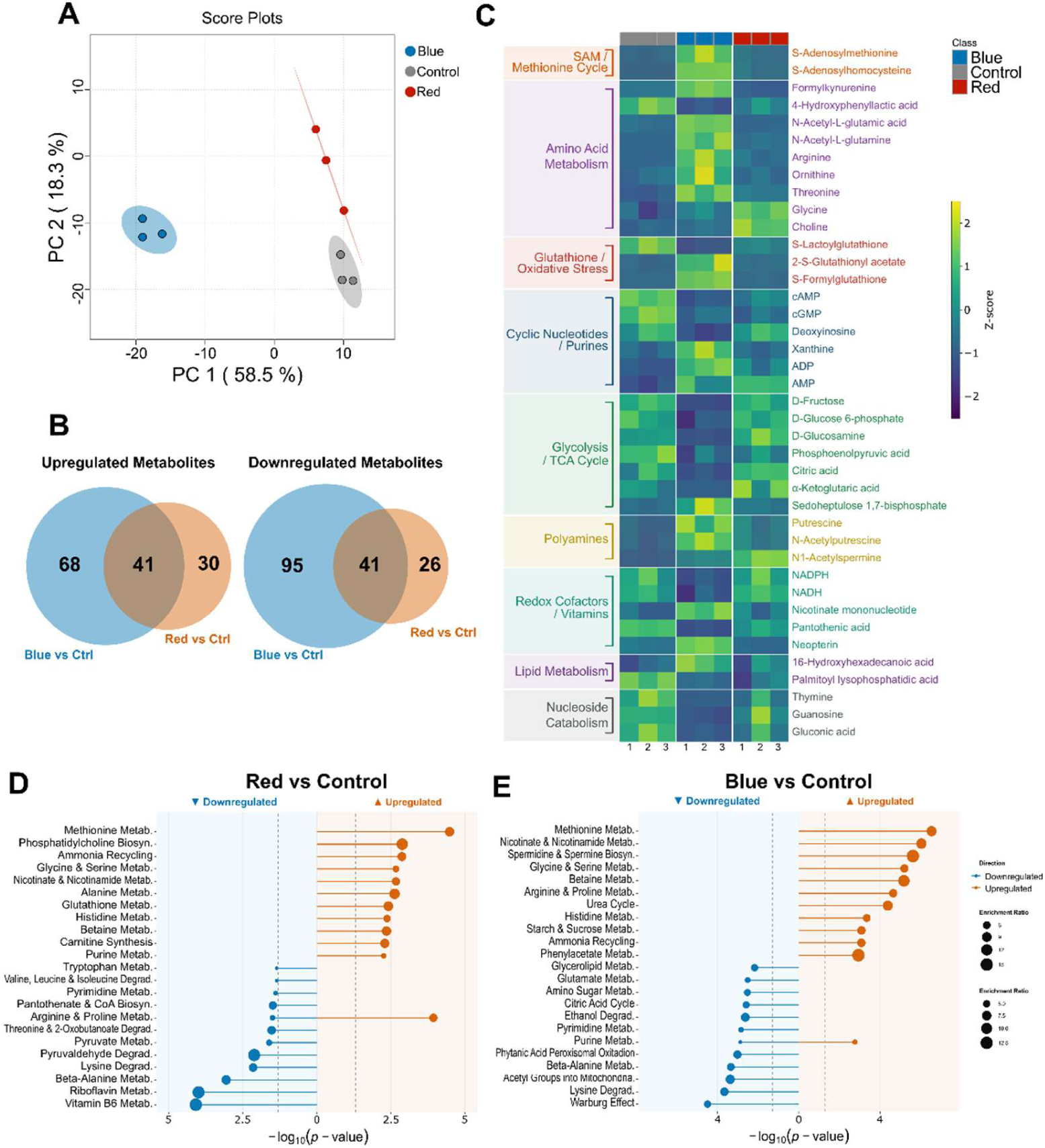
Chromoprotein expression remodels the *E. coli* metabolome in a color-dependent manner. (**A**) PCA score plot of metabolic profiles from HT115 *E. coli* expressing red or blue chromoprotein or no-color control (PC1 = 58.5%, PC2 = 18.3%; *n* = 3 per condition). (**B**) Venn diagrams showing overlap of significantly upregulated (left) and downregulated (right) annotated metabolites between blue vs. control and red vs. control (one-way ANOVA with Fisher’s LSD post-hoc test, FDR < 0.05; n = 3). (**C**) Heatmap of the top 40 differentially abundant annotated metabolites. Color intensity represents Z-score normalized relative abundance; metabolite class annotations are indicated. (**D–E**) Metabolic pathway enrichment analysis for significantly altered metabolites in red vs. control (**D**) and blue vs. control (**E**) bacteria. Bubble size reflects the number of metabolites per pathway; x-axis shows enrichment ratio. Upregulated (▲) and downregulated (▼) pathways are shown separately.

The top differentially abundant metabolites revealed signatures of the methionine cycle, oxidative stress, and redox cofactors (**Fig. 4C**). Formyl kynurenine, a tryptophan catabolite of the kynurenine pathway, was significantly increased in blue bacteria, suggesting diversion of tryptophan away from serotonin synthesis (*24*); this pathway also modulates immune activation and food-related behavioral plasticity in *C. elegans* (*24*). Neopterin, a pteridine related to biopterin, the oxidation product of tetrahydrobiopterin (BH4) (*25*), was likewise altered in blue bacteria, consistent with increased GTP cyclohydrolase I flux and oxidative stress. Neopterin also serves as a cofactor for nitric oxide (NO) synthases, which produce NO. In mammals, inflammation, specifically through the cytokine interferon-gamma, stimulates not only tryptophan catabolism but also the production of neopterin(*25*).

In contrast, red bacteria displayed a metabolite signature consistent with membrane-biosynthetic and amino-acid-handling flux rather than oxidative stress (**Fig. 4C**). Glycine and choline were significantly elevated in red bacteria. Choline contributes directly to phosphatidylcholine synthesis and can be oxidized to betaine, whereas glycine participates in interconnected one-carbon and amino-acid metabolism (*26*), consistent with the significant enrichment of phosphatidylcholine biosynthesis and betaine metabolism among red-upregulated pathways (**Fig. 4D**). α-Ketoglutarate, a central transamination substrate linking the TCA cycle to amino acid and glutathione metabolism, was also increased, aligning with the enrichment of ammonia recycling and glutathione metabolism in red bacteria (**Fig. 4C, D**). Notably, neopterin elevated in blue bacteria as a marker of oxidative, interferon-like activation was decreased in red (**Fig. 4C**), reinforcing a dichotomy between the two chromogenic bacterial food: an oxidative, tryptophan-diverting signature in blue versus a biosynthetic, membrane-and amino-acid-supportive signature in red. Other red-elevated metabolites, such as the polyamine catabolite N1-acetylspermine (**Fig. 4C**), were not accompanied by significant enrichment of the corresponding pathway (e.g., polyamine biosynthesis was enriched in blue, not red). Pathway enrichment corroborated these signatures: methionine metabolism was upregulated in both conditions, with ammonia recycling and phosphatidylcholine biosynthesis in red and urea cycle, spermidine, and betaine metabolism in blue, while downregulated metabolites mapped to pyrimidine, pyruvate, and vitamin B6 metabolism in red and to peroxisomal oxidation, lysine degradation, and citric acid cycle and Warburg pathways in blue, also indicating mitochondrial stress (**Fig. 4D, E**).

RNA-seq of N2 worms fed control, red, or blue *HT115* separated the three diets by PCA of FPKM-normalized data (PC1 = 55.6%, PC2 = 19.1%), mirroring the bacterial metabolomes (**Fig. 5A**). Blue-fed worms displayed 7,035 differentially expressed genes (DEGs; 4,026 up, 3,009 down, *padj ≤ 0.05*) and red-fed worms 6,996 DEGs (2,957 up, 4,039 down), with 1,308 up and 793 down unique to blue and 1,256 up and 801 down unique to red (**Fig. 5B and File S2**).

**Figure 5.**
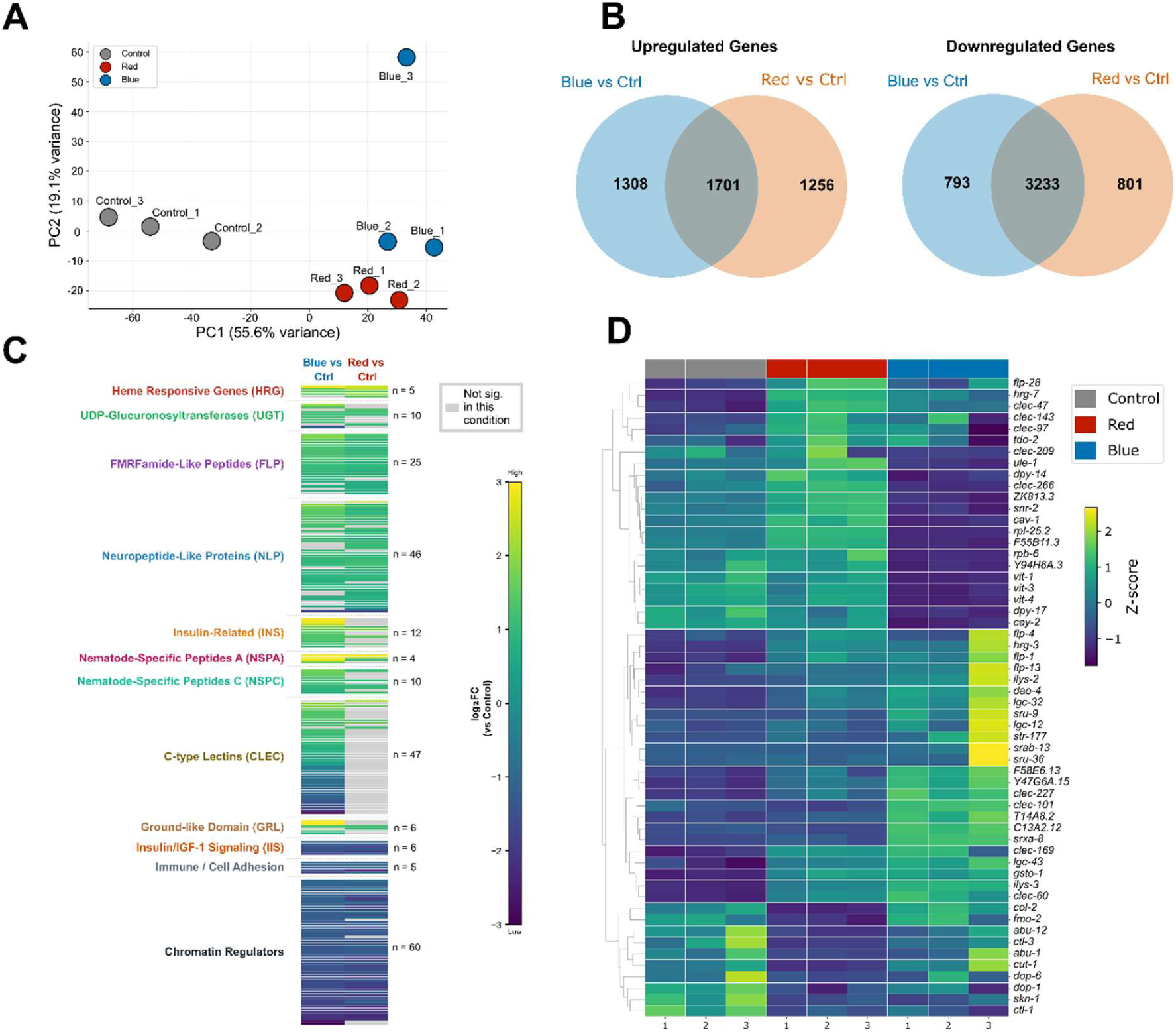
Chromoprotein diet drives divergent transcriptional reprogramming in *C. elegans*. (**A**) PCA of FPKM-normalized gene expression from worms fed no-color control, red, or blue chromoprotein-expressing *E. coli* (PC1 = 55.6%, PC2 = 19.1%; *n* = 3 per condition). (**B**) Venn diagrams showing overlap of upregulated (left) and downregulated (right) DEGs between blue vs. control and red vs. control comparisons (*padj* < 0.05). (**C**) Log₂ fold change plot of curated gene families among significant DEGs (*padj* < 0.05) in blue-fed (left) and red-fed (right) worms relative to control. Each bar represents an individual gene; gray indicates genes not significant in that condition. Gene family sizes are indicated (*n*). (**D**) Hierarchical clustering heatmap of selected DEGs, Z-score normalized across samples. Sample groups are indicated by color bars (gray, Control; red, Red; blue, Blue).

Blue food elicited a broad innate immune and cuticle barrier program consistent with pathogen defence (*27*) . Fifteen C-type lectin genes were significantly upregulated (*padj ≤ 0.05*), including *clec-60*, *clec-227*, and *clec-101*, whereas red-fed animals induced a largely non-overlapping set of five, constituting a lectin repertoire switch between chromatic states (**Fig. 5C, D and File S2**). Blue-fed worms also induced the peptidoglycan-cleaving lysozymes *ilys-3* and *ilys-2* (*28*), 14 cuticle collagens, cuticlin-1 (*cut-1*, log2FC 3.07), the lipid elongase *elo-5*, four ground-like ligands (*grl-20*, *grl-23*, *grd-4*, *grl-25*) reciprocally suppressed, with *grd-11*, under red food, and 23 serpentine/GPCR genes (up to 5.5-fold log₂; *str-177*, *srab-13*, *srxa-8*, *sru-9*, *sru-36*) whose class U and Str receptors are enriched in the aversion-related AWB and ASH neurons (*29*) . Conversely, 13 germline small RNA genes (*wago-1*, *ergo-1*, *pgl-3*, *nrde-3*) and the vitellogenins *vit-1*, *vit-3*, *vit-4*, and *vit-6* were suppressed (**Fig. 5D, File S2)**, and in line with this observed gene expression signature we found that blue-fed worms produced significantly fewer eggs and progeny (*p = 0.0077*, student’s t-test) (**Fig. S14A-C**). Induction of *dao-4* (log₂FC 2.11) (*30*) alongside insulin/IGF-1 signalling suppression further suggests partial entry into a dauer-like state. The strongest signal in the study, *hrg-7* (*padj* = 4.68×10^−77^, log₂FC 2.09), encodes a secreted inter-tissue heme sensor that modulates food-quality behavior under heme depletion (*31*) ; all five *hrg* genes were upregulated in both conditions, indicating heme/iron sequestration by these large β-barrel chromoproteins, most potently by meffRed.

Chromogenic bacteria feeding also drove a coordinated response across the FLP and NLP families: of 31 detected *flp* genes, 22 were regulated under blue and 24 under red, and among NLPs 34 under blue and 43 under red (*padj ≤ 0.05*), almost all upregulated and largely shared (21 of 24 *flp*; 31 *nlp*), spanning motor-neuron and foraging-related interneuron peptides (*flp-1*, *flp-7*, *flp-8*, *flp-9*, *flp-17*, *flp-20*, *flp-22*) and constituting a common signature of chromogenic bacteria exposure rather than a color-specific signal (**Fig. 5D and File S2**). Condition-selective peptides, by contrast, distinguished blue danger-like transcriptional program from red safety. Blue selectively induced *flp-13* (log2FC +0.829, *padj* = 2.04×10^−2^), a mediator of sickness behavior and stress-induced quiescence (*32*), and a *flp-13::GFP* reporter (CZ2475) (*33*) showed significantly increased expression after 24 hours on blue but not red bacteria (*p = 0.0001*) (**Fig.6A, B**).

**Figure 6.**
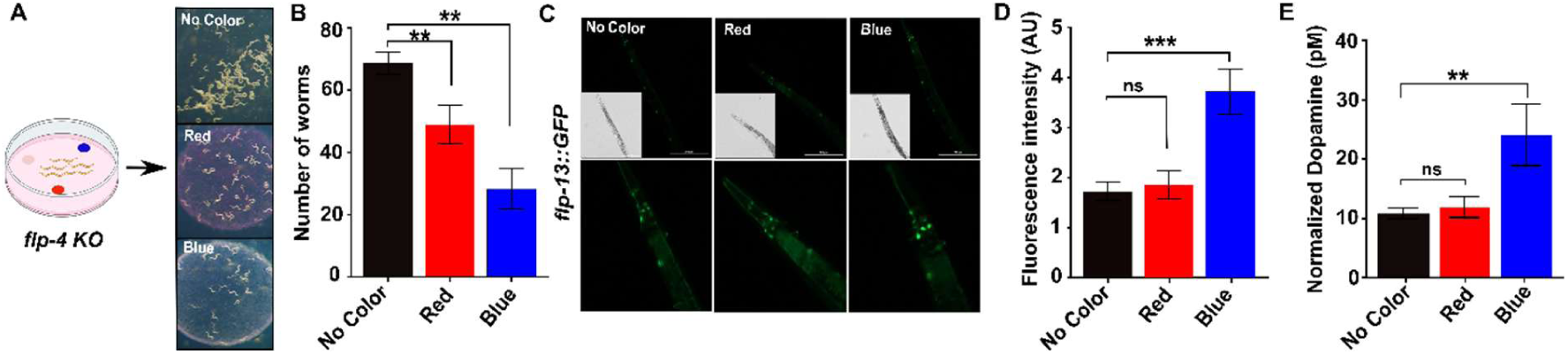
Neuropeptide signaling plays role in chromatic bacterial food discrimination. (**A**) Loss of *flp-4* disrupts red chromoprotein preference in *C. elegans*. Representative images of chromoprotein choice assays performed using *flp-4* knockout worms exposed to non-chromogenic control, red, or blue chromoprotein-expressing bacteria. Synchronized worms were transferred at the L4 stage and allowed to choose among the indicated bacterial lawns for 24 h under ambient light conditions. (**B**) Quantification of bacterial choice in *flp-4* knockout worms after 24 h of exposure. Data are presented as mean ± SEM from three independent biological replicates (*n* = 3). Statistical significance (\**p=0.0144,0.0023*) was determined using an unpaired Student’s *t* test. **(C)** Blue chromoprotein-expressing bacteria increase FLP-13::GFP expression in *C. elegans*. Representative fluorescence images of FLP-13::GFP reporter worms exposed to non-chromogenic control or blue chromoprotein-expressing bacteria. Synchronized worms were transferred at the L4 stage to the indicated bacterial diets and maintained for 24 h under ambient light conditions. (**D**) Quantification of FLP-13::GFP fluorescence intensity following 24 h of exposure. Worms fed blue chromoprotein-expressing bacteria exhibited significantly increased FLP-13::GFP fluorescence compared with control-fed worms. Fluorescence was quantified as the mean fluorescence intensity per worm (*n* = 15 worms per condition). Data are presented as mean ± SEM. Statistical significance was determined using an unpaired Student’s *t* test (*p* < *0.001*). (**E**) The bar graph shows dopamine levels in wild-type (N2) worms following 24 h exposure to control and red and blue chromophore-producing bacteria. Dopamine levels were significantly altered in the blue chromogenic bacteria fed worms (\**p* = 0.048, Student’s *t*-test).

Red food selectively upregulated *flp-4* (log2FC +0.703, *padj* = 1.33×10^−2^), which mediates value-based choice between alternative food cues (*34*), and *flp-21* (+0.506, near-threshold *padj* = 0.051), which mediates acute chemosensory responses (*35*) (**Fig. 6C and File S2**). Accordingly, *flp-4* knockout worms (PS9050) (*36*) lost red attraction while retaining blue avoidance (**Fig. 6D**). Ten NLP genes were upregulated exclusively under red, including the ASI-expressed opioid-like feeding regulator *nlp-24* (+0.737, *padj* = 2.09×10^−6^) (*37*) and the inducible epidermal antimicrobial peptide *nlp-33* (+0.625, *padj* = 3.19×10^−3^) (*38*), whereas *nlp-14*, an ALA neuron neuromodulator governing rest and stress-induced quiescence (*39*), was the sole NLP gene downregulated specifically under red (−0.442, *padj* = 2.11×10^−2^). Red food thus promotes feeding peptides (*nlp-24*, *nlp-63*, *flp-4*) while suppressing quiescence-driving *nlp-14*, the mirror image of the blue-induced *flp-13*, *nlp-46*, and *nlp-12* program. Red feeding also suppressed the dopamine receptors *dop-6* (log2FC −3.71, *padj* = 3.5×10^−4^) and *dop-1* (**Fig. 5D**); because dopamine from mechanosensory neurons mediates aversive food-quality memory through DOP-6 on AWB neurons (*40*), this receptor suppression, combined with neuropeptide activation, constitutes a dual mechanism for attraction. We further validated this observation by measuring dopamine levels using LC-MS/MS. We found that worms fed blue bacteria exhibited significantly increased (*p = 0.048*, student’s t-test) dopamine levels (**Fig. 6E**). In the nematode *C. elegans*, dopamine acts as a key neurosensory “danger” signal, linking environmental stressors, including pathogenic bacteria and physical invasion cues, to cellular stress responses and innate immune defenses (*41*, *42*). This increase in dopamine therefore further aligns with the overall gene expression pattern observed under blue-food conditions.

Together, the gene expression analyses and experimental validation of the transcriptomic findings support a novel model of binary foraging decisions in response to chromogenic bacterial food: blue food promotes the allocation of resources toward defense and stress responses, whereas red food favors neuromodulatory reward signaling and safe chromophore processing.

### Blue chromophore avoidance behavior is mediated through oxidative stress

To validate blue avoidance, we grew N2 worms on no-color bacteria to the young adult stage, transferred them onto blue or red lawns, and calculated retention rate (**Fig. 7A**). Significantly more worms remained on red lawns than left them (*p = 0.005*, student’s t-test), whereas significantly more escaped blue lawns than remained (*p = 0.003*, student’s t-test), relative to no-color control (**Fig. 7B**). Blue-fed worms also showed significantly increased reactive oxygen species (ROS) (*p = 0.0297*), whereas red-fed worms were unchanged (**Fig. 7C**), and blue bacteria were themselves significantly more sensitive to hydrogen peroxide (*p = 0.0151*) (**Fig. 7D, E**), indicating that blue chromophore expression induces intrinsic bacterial oxidative stress, consistent with the oxidative stress-related metabolites detected by metabolomics (**Fig. 4C**).

**Figure 7.**
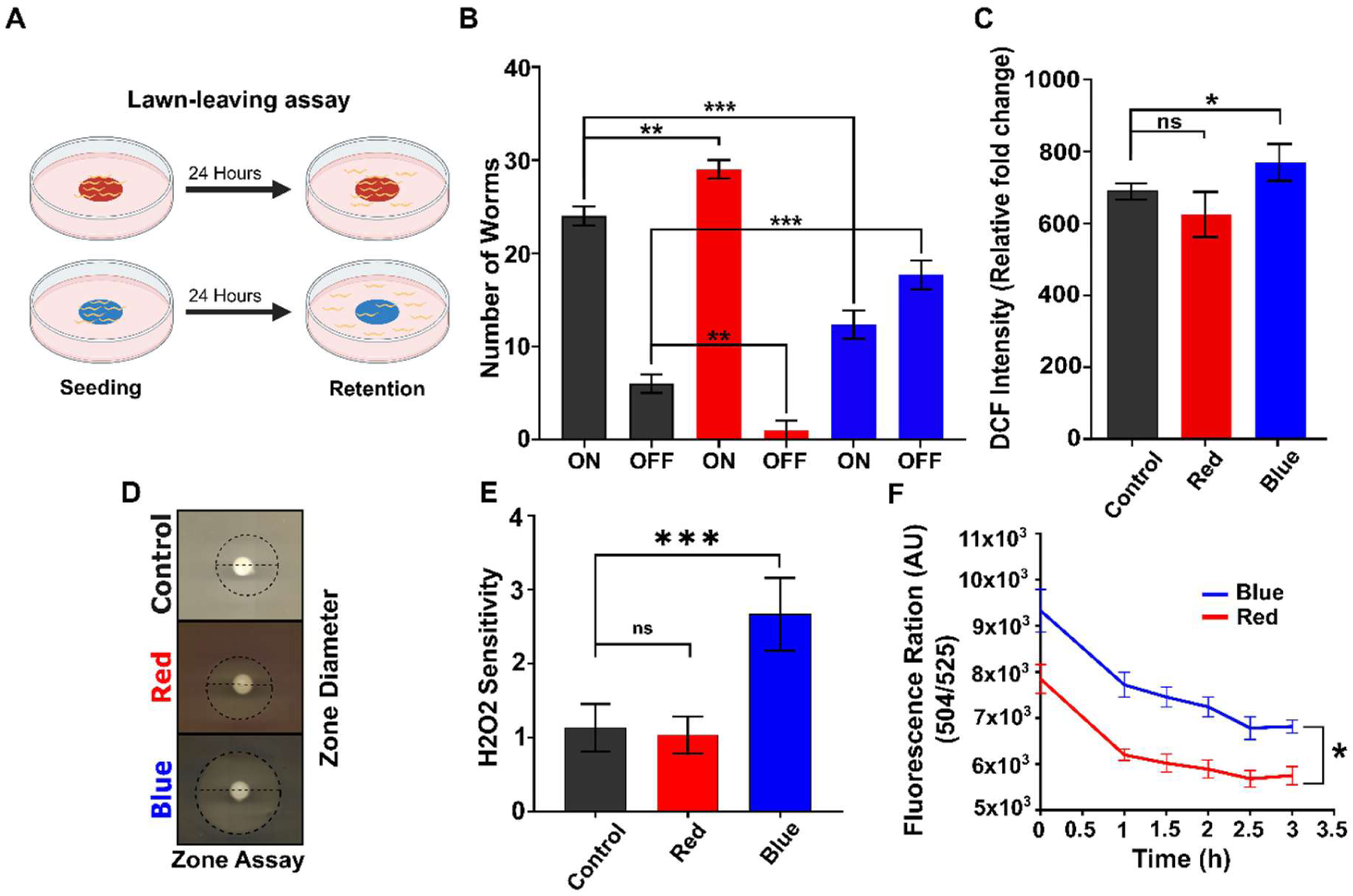
Blue chromophore bacteria induces oxidative stress in in worms and bacterial cells: **(A)** Depiction of experimental procedure for the lawn-leaving behavior assay where young adult worms were placed onto a bacterial lawn of red, blue and no color chromogenic bacteria (HT115) seeded at the canter of NGM plates and allowed to roam freely up to 24 hours. **(B)** Bar graph represents the result from lawn leaving assay showing N2 worms remained on each chromogenic bacterial lawn along with no color control after 24 hours (n=3 and 100 worms on each plate were assayed, \**p = 0.005, 0.003* student’s t-test). **(C)** Bar graph shows the ROS levels at young adult N2 worms reared on no color, red and blue chromogenic bacterial cultures (n=100, *p = 0.0297,* student’s t-test). (**D**) Plate images of bacterial zone assay. Standard LB plates were seeded with red and blue chromogenic bacterial cultures along with no color control and H2O2 sensitivity was measured. (**E**) The bar graph shows the result from the zone assay (n= 3, *p= 0.0151,* student’s t-test). (**F**) Singlet oxygen production was quantified using Singlet Oxygen Sensor Green (SOSG) assay kit for purified red and blue chromophores (n= 3, *p = 0.0265,* student’s t-test). All assays presented in this figure were done on standard NGM plates with HT115 bacteria.

These results do not explain how aeBlue induces oxidative stress or how the purified blue chromophore triggers avoidance independent of bacterial metabolic output (**Fig. 3**). Both aeBlue and aeRed are non-fluorescent chromoproteins with twisted, *trans* non-coplanar chromophore conformations (*43*); because neither fluoresces, absorbed photon energy must dissipate through internal conversion or intersystem crossing to a reactive triplet state (³CP*) capable of Type II photosensitization with molecular oxygen, generating singlet oxygen (¹O₂) (*44*). Accordingly, purified aeBlue in liquid nematode growth medium produced significantly higher ¹O₂ levels than aeRed (*p = 0.0265*) (**Fig. 7F**). Chromophore structure offers an explanation: the aeBlue chromophore (ε = 122,573 M⁻¹cm⁻¹ at ∼608 nm) carries a phenolic group with a pKa of ∼10.46 and remains predominantly protonated (-OH) at physiological pH (*43*); because protonated phenols quench singlet oxygen inefficiently, ¹O₂ escapes into the aqueous phase, establishing a persistently oxidizing microenvironment sensed by worms. aeRed likely has a lower effective pKa, stabilizing the deprotonated phenolate (-O⁻) and a far less oxidizing microenvironment.

Three transcriptional signatures confirm this asymmetric oxidative burden. First, three cys-loop ligand-gated ion channels with regulatory cysteines in their ligand-binding domains were induced exclusively in blue-fed worms, *lgc-32* (log₂FC +1.37, *padj* = 1.15×10⁻⁶), *lgc-43* (+1.13, *padj* = 1.16×10⁻⁴), and *lgc-12* (+2.03, *padj* = 8.76×10⁻³), consistent with selective activation of oxidation-sensitive ionotropic receptors. Second, the methionine sulfoxide reductase *msra-1*, which repairs singlet oxygen-oxidized methionine (*45*), was upregulated specifically in blue-fed worms (+1.15, *padj* = 1.08×10⁻⁹) and absent from the red DEG list (**File S2**); because methionine is a primary ¹O₂ target (*43*, *44*), this indicates a profound oxidative burden under blue food and suggests that quenched ¹O₂ in the aeRed condition fails to reach worm protein targets. Third, the H₂O₂-scavenging catalases *ctl-3* (−2.10, *padj* = 4.33×10⁻¹¹) and *ctl-1* (−1.05, *padj* = 6.11×10⁻⁷) were suppressed exclusively in red-fed worms (**File S2**), reflecting a lower steady-state peroxide burden. Wavelength-selective photosensitization of the blue chromophore therefore elevates ROS in both bacteria and worms, shaping foraging behavior.

### Neuropeptide serotonin circuit drives red color preference in *C. elegans*

The neuropeptide transcriptional changes, together with the loss of red attraction in *flp-4* mutants, led us to hypothesize that a neuroendocrine circuit underlies color-dependent foraging. Using red-versus-blue choice assays, we screened mutants lacking key neurotransmitters: dopamine (*cat-2*: CB1112), glutamate (*eat-4*: MT6308), octopamine (*tbh-1*: RB1161), tyramine (*tdc-1*: MT13113), acetylcholine (*unc-17*: CB3031), GABA (*unc-25*: CB156), and serotonin (*tph-1*: PHX3596) (*46*) (**Table S1**). Most retained the red preference; the exception was the serotonin-deficient *tph-1* line, which foraged on and consumed red and blue lawns equally (**Fig. 8A, B and Fig. S15**). Consistently, the serotonin reporter *tph-1::GFP* (GR1333) showed significantly (*p = 0.004,* student’s t-test) increased expression in red-fed worms (**Fig. 8C, D**), and LC-MS/MS confirmed significantly elevated serotonin relative to worms fed non-chromogenic food *(p = 0.0067*), while there were no alteration in serotonin levels in blue-fed worms (**Fig. 8E, Video S1**).

**Figure 8.**
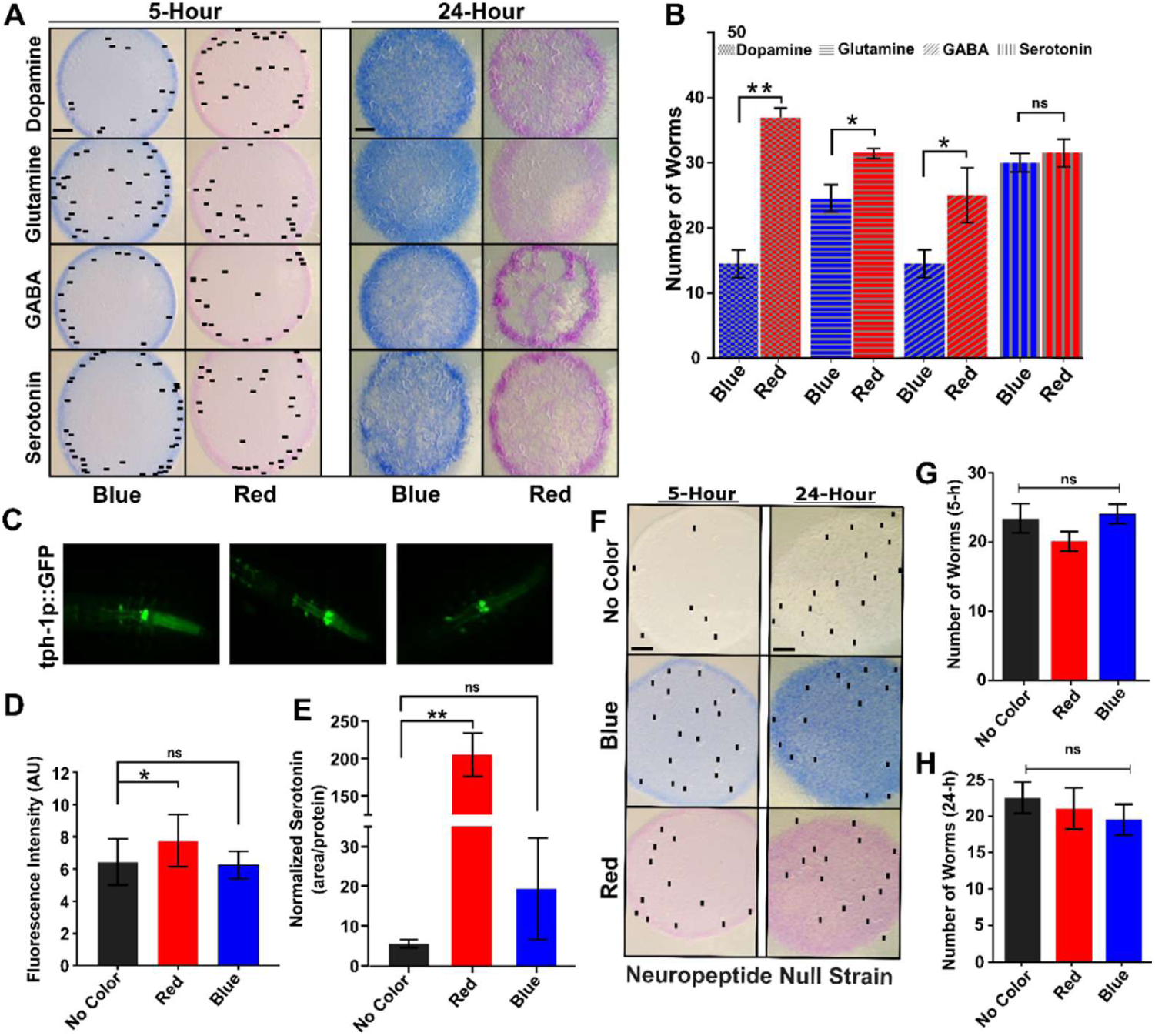
Red chromophore modulates serotonin signaling by regulating neuropeptide expression in *C. elegans*: **(A)** Representative plate images show the chromophore attraction assay of different mutant lines; dopamine (CB1112), Glutamate (MT6308), GABA, (CB156), and Serotonin (PHX3596) after 5-and 24-hours incubation (black dots represent worm locations). **(B)** Bar graph shows the number of worms that were presented on no color (control), red and blue chromogenic bacterial lawn after 5 hours of incubation. (n=3, 50 worms were assayed on each, \**p = 0.05*, student’s t-test). **(C)** Photomicrograph of GR1333 strain (yzIs71 [tph-1p::GFP + rol-6(su1006)] V) reared on no color (control), red and blue chromogenic bacterial cultures. **(D)** Bar graph shows the fluorescence intensity in arbitrary units of GR1333 strain for *tph-1::gfp* expression (n=20, \**p = 0.04*). **(E)** Red chromoprotein-expressing bacteria increase serotonin levels in *C. elegans*. Quantification of serotonin levels in N2 worms following exposure to non-chromogenic control, red, or blue chromoprotein-expressing bacteria. Worms were exposed to the indicated bacterial diets for 24 h, after which serotonin levels were measured by liquid chromatography–mass spectrometry (*p* = *0.0067*; *n* = 3 biological replicates). Data are presented as mean ± SEM. Statistical significance was determined using an unpaired Student’s *t* test. (**F**) Representative plate images show the chromophore choice assay of young adult egl-3 KO worms (MT150) on plates seeded with no color, red and blue chromogenic bacterial lawns. Black dots represent worm locations. Bar graph represents no significant choice of attraction for the chromophores by MT150 worms after (**G**) 5-and (**H)** and 24 hours of incubation (n=100 number of worms assayed on each plate, p > 0.05). The data represent three biological replicates for each experiment.

To test the neuropeptide arm, we examined the neuropeptide-processing mutant MT150 (*egl-3(n150)*) (*47*), which lacks the proprotein convertase required for FLP and NLP maturation. MT150 worms lost both red preference and blue avoidance (**Fig. 8F-H and File S3**), indicating that neuropeptide signalling is required for color-based foraging. Collectively, our data support a model in which chromatic food discrimination operates, after ingestion, through a gut-brain metabolic axis: the ingested chromoprotein’s biochemical properties determine intestinal tryptophan fate, gating serotonin biosynthesis in sensory neurons, which activates neuropeptide circuits that execute the foraging decision, a mode of sensory-to-motor integration mechanistically distinct from previously described chromatic discrimination (**Fig. 9**).

**Figure 9.**
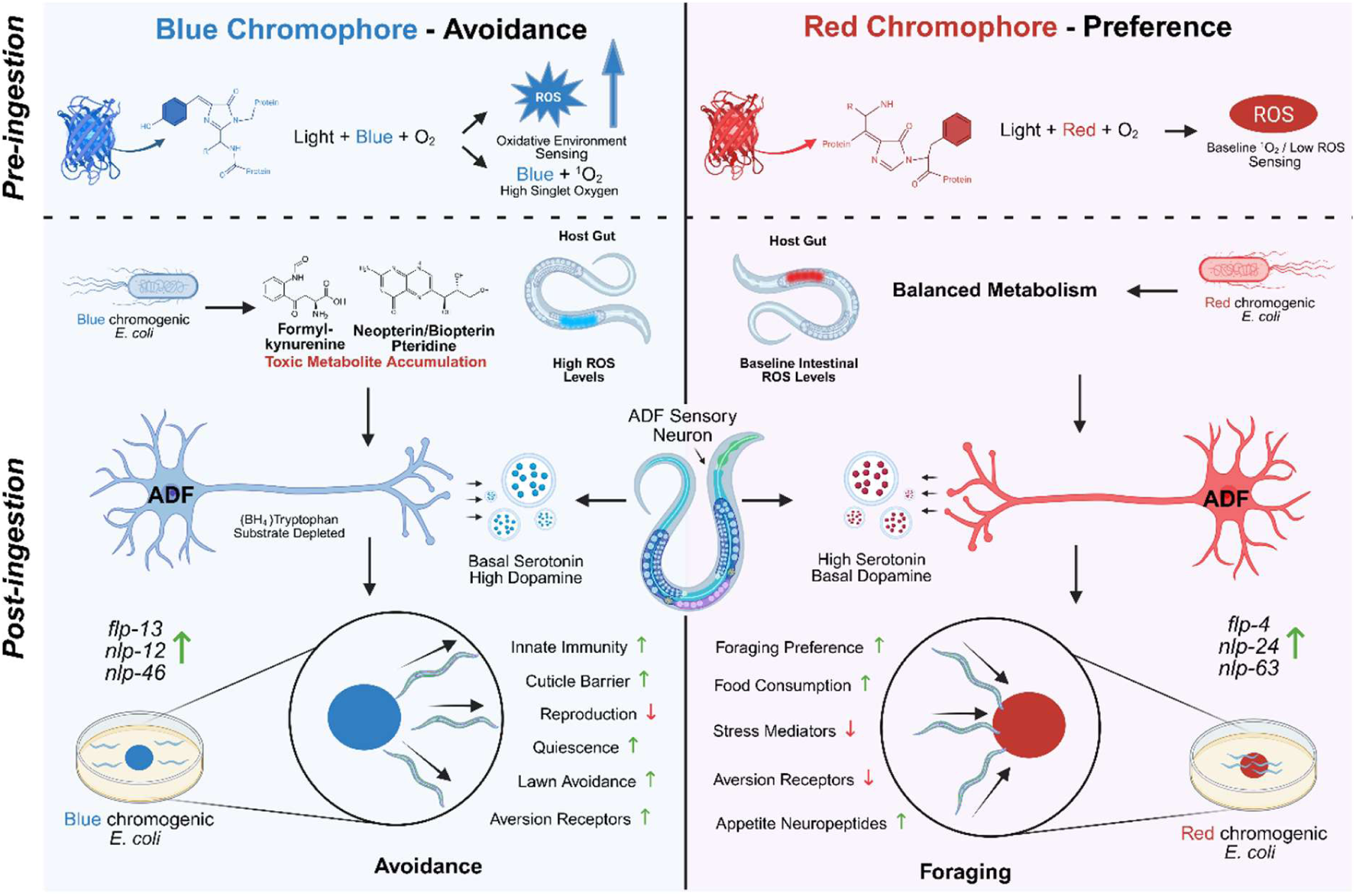
Mechanism of chromatic food discrimination in *C. elegans*. Two discrimination systems for the blue and red-chromophore expressing bacteria are operating in parallel: an initial pre-ingestive system based on wavelength-selective blue chromoprotein photosensitization that resulted in oxidative environment (altered reactive oxygen species [ROS]), mediating an aversion behaviour. The post-ingestive system operates based on intestinal sensing of a pre-formed photochemical metabolite fingerprint that operates first within the bacteria and then within the optically transparent worm body. Both converge on differential neuropeptide synthesis and alter serotonin biosynthesis as the proximate determinant of foraging behavior

## Discussion

Our study reports that *C. elegans* exhibits chromatic food discrimination, robustly preferring red chromophore-expressing bacteria while avoiding blue. Discrimination was fully preserved when worms were presented with a purified His-tagged chromophore devoid of other bacterial components, indicating that the chromophore protein itself is the initial discriminating cue, independent of ingestion, and that a post-ingestive, serotonin-mediated gut-to-neuron mechanism subsequently reinforces the behavior.

Our metabolomic and transcriptomic profiling defines two discrimination systems operating in parallel: an initial pre-ingestive system based on wavelength-selective reactive oxygen species (ROS) detection, and a post-ingestive system based on intestinal sensing of a pre-formed photochemical metabolite fingerprint that operates first within the bacteria and then within the optically transparent worm body. Both converge on differential serotonin biosynthesis as the proximate determinant of foraging behavior.

The first system operates before ingestion, chromatic discrimination may occur through the worm’s nose via the amphid ASH and ADF neurons. Our data indicate that the chromophore deprotonation state dictates the local ROS burden: aeBlue creates a persistently oxidizing microenvironment, while meffRed creates a reducing or neutral one. This asymmetric oxidative load establishes the basis of avoidance of the purified blue chromophore and validates that the chromoprotein alone is sufficient to drive the initial foraging decision. We propose that this extracellular oxidative asymmetry is plausibly transduced at the amphid sensilla via TRPA-1 and OSM-9, constitutively expressed TRP channels on ASH nociceptor neurons whose N-terminal ankyrin cysteines are directly gated by oxidation from singlet oxygen and peroxide, an avoidance mechanism entirely independent of LITE-1. Consistently, the oxidation-sensitive cys-loop channels *lgc-32*, *lgc-43*, and *lgc-12* were induced exclusively in blue-fed worms. Simultaneously, the red chromophore engages ADF serotonergic neurons and bidirectional neuropeptide regulation, with quiescence peptides induced and foraging peptides suppressed under blue food and the reverse under red, a circuit-level switch that amplifies and sustains chromatic discrimination beyond the initial sensory detection event.

The second system, post-ingestive metabolite fingerprint sensing in the intestine, acts through two photosensitization sites. The bacterial metabolome establishes the first. Blue chromophore-expressing bacteria accumulated N-formylkynurenine at markedly elevated levels relative to red bacteria, in which this metabolite was not significantly detected. N-Formylkynurenine is a characteristic product of singlet-oxygen-mediated tryptophan oxidation, a Type II photosensitization reaction that cannot occur by thermal chemistry at physiological conditions and requires ¹O₂ generated from an excited photosensitizer (*45*). aeBlue absorbs ambient ∼608 nm light and generates abundant ¹O₂, which would oxidize bacterial tryptophan to formyl kynurenine and simultaneously photo-oxidize tetrahydrobiopterin (BH4), the electron-donating cofactor of tryptophan hydroxylase-1 (TPH-1), to the pteridine catabolites neopterin and biopterin, both of which accumulated specifically in blue-expressing bacteria. The resulting singlet oxygen may also drive flavin (FAD, FMN, riboflavin) oxidation, producing superoxide and H₂O₂ under ambient light (*48*), consistent with the H₂O₂ sensitivity of blue bacteria. The red chromophore transmits >610 nm, does not excite flavins at this wavelength, and presents an essentially unperturbed metabolome. Blue chromophore-expressing bacteria therefore autonomously convert their absorption wavelength into a distinctive metabolite fingerprint, depleted tryptophan, BH4 and serotonin with accumulated formyl kynurenine, neopterin, and other oxidized metabolites, encoded in the bacterial cytoplasm before the worm first contacts the food.

*C. elegans* is optically transparent, a property that creates a second photosensitization site inside the worm during daylight feeding. When blue bacteria are ingested, the chromophore in the gut lumen continues to absorb ambient photons at its ∼608 nm peak, populates the triplet state, and generates singlet oxygen precisely at the interface with the gut epithelium. These photons may propagate through the gut wall and oxidize the worm’s own intracellular flavins in gut cells and adjacent tissues, extending the tryptophan depletion and BH4 photooxidation cascade into the worm’s own cellular environment.

The two sites act additively on the same molecular target: the tryptophan/BH4-dependent serotonin biosynthetic axis in ADF amphid serotonergic neurons. Site 1 delivers a pre-formed supply of formyl kynurenine (depleting tryptophan substrate) and pteridine catabolites (depleting BH4 cofactor) through intestinal absorption, while Site 2 generates the same depletions in real time within the worm’s gut cells. Upregulation of *ptps-1* (6-pyruvoyl tetrahydrobiopterin synthase, de novo BH4 synthesis) in blue fed condition reflects attempted but insufficient restoration of a BH4 pool depleted from two convergent sources; BH4 is the essential cofactor for serotonin synthesis by TPH-1 (*25*) . Under red feeding, the absence of photosensitization at either site preserves tryptophan and BH4 for full TPH-1 activity, elevating ADF serotonin output as required for red preference. Discrimination in complete darkness is consistent with this dual-site model: Site 2 requires ambient light and is absent in darkness, but Site 1 is light-history dependent, since bacteria grown under standard laboratory lighting accumulate H₂O₂, formyl kynurenine, and neopterin before the behavioral experiment begins, so feeding these pre-loaded bacteria in darkness still delivers a metabolite fingerprint that initiates oxidative stress and impairs serotonin synthesis. Complete LITE-1 independence is likewise explained: neither Site 1 nor Site 2 involves LITE-1, which detects external photons entering through the body wall rather than photons absorbed by ingested chromoproteins in the gut. LITE-1 may instead have evolved for UV/blue-light avoidance from sunlight and may contribute to detecting *P. aeruginosa* pyocyanin in natural pathogen encounters, where blue color is accompanied by real toxins (*8*), whereas the chromophore-expressing *E. coli* used here are otherwise normal food.

Overall, we identify dual-site wavelength-selective photosensitization, operating first in the bacteria as a Photochemical wavelength encoder and then in the transparent worm as a real-time color amplifier, as the primary mechanism by which *C. elegans* transduces chromatic food identity into graded serotonin output and, through neuropeptide signaling, into the decision to forage or avoid. In ambient light, aeBlue generates unquenched singlet oxygen that creates an oxidizing amphid microenvironment driving acute avoidance, while in bacteria the same photosensitization pre-loads the food with formyl kynurenine and neopterin that chronically suppress serotonin synthesis, sustaining avoidance even in darkness. meffRed, whose deprotonated phenolate chromophore quenches its own singlet oxygen, avoids this oxidative signature entirely and presents a reducing, serotonin-permissive environment at both the amphid surface and the intestinal lumen.

These findings reveal an unexpected sensory modality through which *C. elegans* integrates environmental color cues with internal neuroendocrine signaling. In natural habitats such as decomposing organic matter, bacteria display a wide spectrum of pigments, and discriminating among these signals may allow worms to identify beneficial versus potentially harmful food sources; blue pigments often originate from redox-active compounds associated with microbial competition and toxicity, whereas other pigments may signal nutritionally favourable environments. Color-guided foraging may therefore represent an adaptive strategy for navigating complex microbial ecosystems. More broadly, our work demonstrates that even organisms lacking conventional visual systems can exploit environmental color cues to guide behavior: rather than relying on classical photoreceptors, *C. elegans* interprets color-associated chemical and physiological signals through neuroendocrine pathways involving serotonin and neuropeptides. These findings highlight the flexibility of sensory systems and suggest that chromatic information may play a more widespread role in microbial-animal interactions than previously appreciated.

## Materials and Methods

### C. elegans Culture

*C. elegans* strains were cultured on nematode growth medium (NGM) agar plates (90 mm Petri dishes) seeded with *E. coli* OP50 as the food source, and the worms were maintained at 22 °C (*49*). Unless otherwise specified, experiments were performed using plates seeded with live *OP50* or *HT115* bacterial strains. All *C. elegans* strains used in this study were obtained from the Caenorhabditis Genetics Center (CGC), University of Minnesota. A complete list of strains and their genetic backgrounds is provided in (**Table S1**). To generate age-synchronized populations, gravid adult worms were treated with 4-6% sodium hypochlorite (Sigma, Cat: 419550010) to dissolve adult bodies and release embryos (46). The isolated embryos were washed thoroughly with sterile M9 buffer and allowed to hatch and develop under standard growth conditions until they reached the desired developmental stage for each experiment.

### Preparation of Chromogenic Bacterial Strains

To study the color preference in *C. elegans*, we used constructs expressing chromoproteins that are derived from coral reefs (*19*), including, meffRed, eforRed, meffBlue, aeBlue, tsPurple, fwYellow, fuGFP, and scOrange, (**Table S2**). For chromoprotein expression, different *E. coli* strains, including *BL21*, *DH5α*, and *HT115* were transformed with these expression constructs. The transformed bacterial cells were grown on LB Kanamycin plates (50 µg/mL) except meffRed and meffBlue, which were grown on LB chloramphenicol agar plates (30 µg/mL) for 18 hours at 37 °C. Selected bacterial colonies were grown for 36 hours at 37 °C with shaking in LB and the respective antibiotic to express the different chromophores. Visualization of all the chromophores was done in ambient light.

### *C. elegans* Chromogenic Food Choice Assay

Synchronized populations of different *C. elegans* strains were grown on OP50 bacterial food lawns until the young adult stage (*50*). On assay day, worms were washed with M9 buffer three times and once with PBS, followed by the transfer of 100 adult hermaphrodites to the center of the chromophore choice assay plate. These chromophore choice assay plates were prepared 1 day before the experiment. We followed the standard method to design the chromophore choice assay plates, we seeded 25-20ul of 10X live chromogenic bacteria with equal distance on 90mm petri plates or in some cases 60 mm NGM (no antibiotics) plates. These plates were incubated at 22 °C for 24 hours in order to dry the bacterial colonies so that colors are visible differentially. Behavior of different strains of worms towards chromogenic bacterial lawns was recorded/ photomicrographs were taken using a stereoscope (MLJ100 Microscope Digital Camera) at different time points, particularly after 5, 12, and 24 hours, or, in some cases, we followed up for 2 days. We also tracked worms in dark/no-light conditions using the same approach. To quantify the number of worms on respective chromogenic bacterial lawns, we used the established method (*50*) by manually counting the number of worms on each chromogenic bacterial lawn. We plotted these numbers and analyzed the significance using a Student’s t-test.

### Phenotypic Analysis

Behavioral assays were performed using age-synchronized young adult worms (N2-wild type). Pharyngeal pumping was assessed as a measure of feeding behavior by manually counting terminal bulb contractions under a stereomicroscope (Zeiss, 40× magnification) for 30 s at room temperature on different chromogenic bacteria (*51*). Locomotor activity was evaluated using a thrashing assay. Individual worms were transferred to a drop of M9 buffer on a glass slide and allowed to acclimate briefly. A thrashing rate was defined as one complete change in the direction of body bending. Thrashes were counted manually for 30 s. For each condition, 15–20 worms were analyzed per biological replicate, and all assays were performed in three independent biological replicates. Data are presented as the mean ± SEM.

### Lawn Avoidance Assay

Lawn avoidance assays were performed to determine whether *C. elegans* recognizes and avoids chromogenic bacterial lawns as described previously (*52*). Here, we used synchronized young adult hermaphrodite worms that were washed off from NGM plates with M9 buffer, washed thrice, and allowed to settle by gravity. We examined 100 worms in each group, which were placed on top of the bacterial lawn. Plates were incubated for 24 hours at 22 °C. Worms on the bacterial lawn are non-avoiders, called ON lawn worms. Worms that avoid the lawn are named OFF worms. Quantification of the lawn avoidance assay was performed by directly counting the number of worms ON and OFF on the chromogenic bacterial lawn. Data analyses were done using GraphPad Prism.

### Column Separation of Total Bacterial Metabolite and Protein

We developed a centrifugation-based method to fractionate the total protein and total metabolites from chromogenic total bacterial lysate of HT115 strains of *E. coli*. *HT115* chromogenic bacterial colonies were grown overnight, and the next day we reinoculated into a secondary culture at 37 °C in LB medium with kanamycin (50 µg/mL). Next, chromogenic bacterial pellets were harvested by centrifugation at 3500 rpm, 15 min, at 4 °C. The bacterial pellets were washed twice with cold 1X PBS (pH 7.4). Sonication was performed for 3 minutes in 10 second bursts, with 30 second intervals, at 25% amplitude on ice. Resulted lysates were centrifuged at 15,000 × g for 20 min at 4 °C to remove debris and to obtain the total bacterial lysates. To fractionate these total lysates into total metabolite and protein fractions including chromoproteins, we used an ultrafiltration column with a 10 kDa filter (GE Healthcare; 10kDa MWCO-28-9322-25). Proteins ≥10 kDa are retained in the upper part, and small molecules/metabolites pass through the filter to form the lower fraction. Centrifugation of the loaded column was done at 14,000 × g for 30 min at 4 °C. The fractionated total proteins mix was washed three times with 1X PBS before using in our experiments.

### Protein Purification

His-tagged chromophore proteins were expressed in the *E. coli* HT115 strain. A single colony from transformed bacteria expressing chromoproteins was grown overnight at 37 °C in LB medium containing kanamycin (50 µg/mL). This overnight culture was used as the primary culture and diluted 1:1000 into fresh LB containing kanamycin, then shaken at 37 °C for 36 hours. Chromogenic bacterial cells were harvested by centrifugation at 3500 rpm for 20 minutes at 4 °C. The bacterial pellet was washed 3 times with 1X PBS (pH 7.4), then resuspended in bacterial cell lysis buffer (GOLDBIO, cat: GB-177-100). This bacterial suspension was incubated at 37 °C for 30 minutes. At the end of incubation, this suspension was vortexed for 30 seconds to complete the lysis. After the lysis, EDTA was added at a final concentration of 2.5 mM. To completely lyse the bacterial cells, sonication was performed for 3 minutes, with 10 s bursts and 30 s intervals at 25 amplitudes on ice between cycles. This colored bacterial lysate was centrifuged at 15,000 × g for 30 min at 4 °C to remove debris. The total supernatant was loaded onto the complete His-tag purification column (Sigma-Aldrich; version 03; 06781543001). After binding, this column was washed with 5 ml of wash buffer (50 mM Tris-HCl, pH 8.0, 300 mM NaCl, 20 mM imidazole, 0.1 mM EDTA, and 1mM fresh PMSF). After elution, we used the elution buffer (50 mM Tris-HCl, pH 8.0, 50 mM NaCl, 300 mM imidazole, 0.1 mM EDTA, and 1 mM fresh PMSF). The colored eluates containing chromoproteins were collected and passed through an ultrafiltration column with a 10 kDa filter (GE Healthcare; 10kDa MWCO-28-9322-25) to obtain pure chromoproteins and remove excess buffer and salts. These pure chromoproteins were washed 3 times with 1X PBS and once with pure water and were then used as pure color proteins in our experiments.

### RNA-Seq and Data Analyses

*C. elegans* wild-type N2 animals were maintained on NGM plates seeded with *E. coli* HT115 at 22°C following standard protocols. For chromophore feeding experiments, *HT115* bacteria expressing either a blue chromophore or a red chromophore were used. An *HT115* strain carrying the empty vector served as the control. Worms were synchronized by hypochlorite treatment and fed with the respective bacterial strains until reaching young adult stage (L4). Worms were then harvested by washing plates with M9 buffer, and the collected suspension was centrifuged at 2500 rpm for 5 minutes to pellet the animals. The worm pellets were washed three times with sterile M9 buffer to remove residual bacteria and debris, followed by a single wash with 1X PBS to reduce salt contamination. After the final wash, the supernatant was carefully removed, and the worm pellet was immediately flash-frozen in liquid nitrogen.

Total RNA was isolated using column-based kit (Zymo Research, Cat: R2072). Three biological replicates were used per group. RNA integrity was assessed using the Bioanalyzer 2100 system (Agilent Technologies, CA, USA). Strand-specific mRNA libraries were prepared by purifying messenger RNA from total RNA using poly-T oligo-attached magnetic beads. Following fragmentation, first-strand cDNA was synthesized using random hexamer primers, and second-strand cDNA was synthesized using dUTP in place of dTTP to preserve strand directionality. Libraries were finalized through end repair, A-tailing, adapter ligation, size selection, amplification, and purification, then validated by Qubit fluorometry, real-time PCR, and Bioanalyzer. Libraries were sequenced on an Illumina NovaSeq 6000 platform using paired-end chemistry by Novogene Co., Ltd.

For data analysis, raw reads in FASTQ format were quality-filtered using fastp to remove adapter sequences, poly-N-containing reads, and low-quality reads. Quality metrics (Q20, Q30, GC content) were calculated for all retained clean reads. Clean reads were aligned to the *C. elegans* reference genome (NCBI GCF_000002985.6, WBcel235) using HISAT2 (v2.0.5), which generates a splice-junction database from gene model annotations to improve alignment accuracy. Mapped reads were assembled into transcripts using StringTie (v1.3.3b) and read counts per gene were quantified using featureCounts (v1.5.0-p3). Gene expression levels were reported as FPKM (Fragments Per Kilobase of transcript per Million mapped reads).

Differential expression analysis was performed separately for blue chromophore-fed vs. control and red chromophore-fed vs. control comparisons using the DESeq2 R package (v1.20.0), which models count data using a negative binomial distribution. P-values were adjusted for multiple testing using the Benjamini-Hochberg method, and genes with an adjusted p ≤ 0.05 were considered significantly differentially expressed.

### Fecundity Assay

Fecundity was assessed by measuring the number of progeny produced by individual *Caenorhabditis elegans* hermaphrodites. Synchronized worms were maintained on NGM plates seeded with OP50 bacteria under standard culture conditions. At the L4 stage, individual worms were transferred to fresh NGM plates seeded with chromophore-producing bacteria and maintained at 22°C. Each adult worm was transferred individually to a fresh plate every 24 h throughout the reproductive period to prevent overlap between generations. The number of eggs or embryos laid by each worm during each 24-h period was counted using a stereomicroscope (AmScope). Following hatching, the resulting progeny were also counted to determine total offspring production per individual. Fecundity was expressed as the number of viable offspring produced per worm per day. At least 10 individual worms were analyzed per experimental group, and all treatment groups were maintained and assessed under identical experimental conditions.

### Untargeted Metabolomics and High-Resolution LC–MS Profiling

Chromogenic bacteria were prepared for untargeted metabolomics as described (*53*)(*43*). A single colony of each engineered *E. coli* HT115 strain was inoculated into 5 mL LB with kanamycin (50 µg/mL) and grown overnight at 37 °C with shaking for 16 h. Secondary cultures were seeded the following day at 1:10 into fresh LB and grown for a further 36 h under the same conditions. Cells were harvested (4,000 × g, 20 min) and washed twice in sterile 1X PBS. To approximate the nutritional and physiological conditions of NGM plates, pellets were resuspended in 100 mL liquid NGM (without agar) and incubated for 3 h at 22 °C. Bacteria were then recollected under the same centrifugation conditions, washed three times in 1X PBS, and rinsed once in 10 mL of 150 mM ammonium acetate, after which the supernatant was discarded. Pellet volumes were normalized across samples (∼200 µL per sample; three biological replicates). Samples were snap-frozen in liquid nitrogen and stored at −80 °C until extraction.

Frozen samples were thawed on ice, and 874 µL of ice-cold methanol/dH₂O (400:85, v/v) was added to each, followed by sonication to lyse the cells. A further 540 µL of ice-cold chloroform was added, with a second round of sonication where required. Then 360 µL of ice-cold chloroform and 360 µL of ice-cold dH₂O were added to the lysate. Samples were vortexed for 10–15 min at 4 °C and centrifuged at 10,000 × g for 10 min at 4 °C, and the upper polar phase was collected. Extracts were dried in a refrigerated SpeedVac concentrator and stored at −80 °C until analysis.

Dried polar metabolites were reconstituted in 150 µL Optima-grade water. A pooled quality control (QC) sample was prepared from equal volumes of all samples and used to condition the column and to monitor internal-standard intensity, with QC injections after every 10 samples. Ten µL of each reconstituted sample was injected into a Vanquish ultra-high-performance liquid chromatography (UHPLC) system coupled to a Q Exactive HF mass spectrometer (Thermo Fisher Scientific, Waltham, MA, USA) via a heated electrospray ionization (HESI-II) source operating in positive (+ESI) and negative (−ESI) modes. Metabolites were separated on dual Acquity high-strength silica (HSS) pentafluorophenyl (PFP) columns (150 mm × 2.1 mm, 1.8 µm; Waters) at 30 °C and a flow rate of 500 µL/min. Mobile phase A was 0.1% formic acid in water (v/v) and mobile phase B was 0.1% formic acid in acetonitrile (v/v). A 15-min gradient was applied as follows (%A): 98% from 0–3.5 min, 75% from 3.5–11.5 min, 5% from 11.5–12.5 min, and 5% from 12.5–15.0 min.

MS1 data were collected in +ESI and −ESI modes with the following parameters: m/z scanned from 60 to 800 at a resolution of 120,000 at m/z 200; automatic gain control (AGC) target 1 × 10⁶; maximum injection time 100 ms; probe heater temperature 160 °C; sheath gas 53; auxiliary gas 14; sweep gas 10; capillary temperature 320 °C; and S-lens RF level 50. Spray voltages were 3.5 kV and 3.0 kV for positive and negative modes, respectively.

Data-dependent fragmentation (ddMS²) was performed on the five most abundant precursor ions from each full MS scan, in both ionization modes, for a representative sample of each class and for the pooled QC. Parameters differed from MS1 as follows: resolution 30,000; AGC target 1 × 10⁵; maximum injection time 50 ms; and higher-energy collisional dissociation (HCD) at a normalized collision energy of 50 with 50% stepped collision energy. LC–MS data were processed with the XCMS R package for peak detection, mass spectral deconvolution, retention-time alignment, and feature grouping. Mass tolerance was set to 5 ppm and retention-time tolerance to 0.2 min. Features from both ionization modes were filtered to exclude those eluting within the void volume. Background and solvent-derived ions were manually curated and removed on the basis of their average intensity in processing blanks and their coefficient of variation. Metabolites were annotated in Compound Discoverer 3.3 (Thermo Fisher Scientific) using the ddMS² spectra. Features from positive and negative modes were combined, and duplicate annotations were removed on the basis of confidence score and singleton detection in the pooled QC samples. Retained feature intensities were normalized using the MS total useful signal (MSTUS) approach prior to statistical analysis.

Principal component analysis (PCA) was performed in MetaboAnalyst 6.0. Pairwise comparisons of normalized annotated metabolites were performed using one-way ANOVA followed by Fisher’s LSD post-hoc test with false discovery rate (FDR) correction, and adjusted *P* < 0.05 was considered significant. Hierarchical clustering and heatmaps were generated in R v4.2. Metabolite enrichment analysis was performed in MetaboAnalyst 6.0 using qualitative enrichment analysis against KEGG pathway sets.

### LC–MS Measurement of Serotonin

Sample preparation for serotonin analysis was performed according to the protocol described by Schumacher et al (*54*). Wild-type (N2) *C. elegans* were exposed to chromogenic bacteria for 24 h. Following exposure, worms were collected and washed three times with M9 buffer to remove residual bacteria. Approximately 300 mg of worm pellet was collected from each sample and used for total protein/metabolite extraction. The worm pellet was then resuspended in 300 µL of extraction buffer containing 0.1 M perchloric acid and 40 mM sodium thiosulfate. The samples were sonicated for 3 min using a 30 s ON/30 s OFF cycle and subsequently centrifuged at 13,000 × g for 30 min. The resulting supernatants were collected and processed for subsequent quantification of serotonin.

Protein concentration was measured using a NanoDrop spectrophotometer for subsequent normalization of metabolite levels. The total extract was evaporated to dryness using a SpeedVac concentrator and reconstituted in 40 µL of LC–MS-grade water (LiChrosolv™; MilliporeSigma, Cat. No. 01-100-250). Following a 10 second brief centrifugation, the supernatant was collected and used for LC–MS analysis (*55*). For each sample, 5 µL of the 40 µL reconstituted extract was injected into the LC–MS system with two blanks. Liquid Chromatography Chromatographic separation was performed using a Shimadzu Nexera LC-40 XR UHPLC system coupled to a SCIEX X500R quadrupole time-of-flight (QTOF) mass spectrometer. Separation was performed using a Kinetex Polar C18 column (2.6 µm; Phenomenex; Cat. No. 00F-4759-AN) maintained at 30 °C. Samples were analyzed in randomized order. After 10 min equilibration, three pre-run blanks were injected to stabilize and clean the LC column. A serial dilution of a serotonin standard prepared in LC–MS-grade water was also analyzed (serotonin hydrochloride, ≥98% purity; Cayman Chemical, Cat. No. 14332).

Serotonin was identified by comparison of its retention time with that of the authentic serotonin standard and by comparison with the NIST 17 spectral library, providing Level 1 identification confidence. Peak areas for serotonin were determined using SCIEX OS software (version 3.3.1), with a peak-height threshold of 10 arbitrary units and Gaussian smoothing over 10 scans. Absolute serotonin concentrations in the samples were calculated using a calibration curve generated from serially diluted serotonin standards. Protein concentration measured before by Nanodrop is used to normalized per protein metabolite level.

### Measurement of Reactive Oxygen Species

Reactive oxygen species (ROS) levels were measured as per standard procedure using 2′,7’-dichlorodihydrofluorescein diacetate (Sigma, Cat. No. D6883) (*56*). Wild-type (N2) Worms were cultured on chromogenic bacteria, washed 3 times with M9 buffer and once with 1X PBS. For each group, 100 worms were suspended in 100 μl of assay solution and analyzed in triplicate. Worm suspensions were then transferred into wells of a 96-well plate. Fluorescence intensity was recorded at three timepoints: (i) prior to dye addition, (ii) immediately after dye addition, and (iii) after 1 h of incubation. Measurements were obtained using a multimode plate reader (Agilent C10) with excitation at 485 nm and emission at 520 nm. The net fluorescence changes were calculated by subtracting the baseline (pre-dye) reading from the immediate post-dye signal, and this value was further subtracted from the 1 h post-fluorescence. Fluorescence intensity per worm was expressed as the mean value across replicates. Statistical significance was assessed using Student’s t-test in GraphPad Prism software.

### Measurement of Singlet Oxygen

Singlet oxygen production was measured using Singlet Oxygen Sensor Green (SOSG). SOSG was prepared according to the manufacturer’s instructions and used at a final concentration of 2 µM in phosphate-buffered saline (PBS)(*57*). Purified chromophore proteins were added at a final concentration of 10 µM in a total volume of 100 µL in black 96-well plates. Samples were incubated in the dark for 5 min, and baseline fluorescence was measured using a microplate reader (excitation 488–504 nm, emission 525 nm). The plates were then exposed to ambient laboratory light for 1, 2, or 3 h, and fluorescence was measured again using the same settings. Singlet oxygen production was calculated as the change in fluorescence relative to the initial reading: ΔF/F₀ = (F_light − F_dark)/F_dark. Control reactions without chromophore protein or SOSG were included to measure background fluorescence. All experiments were performed in three independent biological replicates, and data are presented as mean ± SEM.

### Hydrogen Peroxide Sensitivity Assay

Overnight bacterial cultures were grown in LB broth at 37 °C with shaking (180 rpm). The following day, cultures were diluted into fresh LB medium and grown to mid-log phase (OD₆₀₀ = 0.5–0.8). To evaluate oxidative-stress sensitivity, bacterial cultures (Red, Blue, and No color-*HT115*) were seeded on an LB agar plate. Subsequently, 6 µL of 30% H₂O₂ (Sigma, Cat: H1009) was spotted onto a sterile paper disc placed on the seeded LB agar plates and allowed to air dry under sterile conditions. Plates were incubated overnight at 37 °C, and the bacterial growth zone was documented by imaging the plates the following day. Oxidative-stress sensitivity was determined by comparing growth patterns and by the presence of a zone of inhibition around the H₂O₂ spot in the control and H₂O₂-treated bacteria. Analysis was performed by measuring the zone dimensions around the H₂O₂ spots across the samples.

### Photomicrographs

Photomicrographs of *C. elegans* strains were taken at the young adult stage. Synchronized worms were washed three times with M9 buffer, once with 1X PBS, and subsequently immobilized using 100 mM sodium azide (Sigma, Cat. No. 71289) (*58*). Immobilized worms were mounted on glass slides and under coverslips to ensure uniform flattening. Fluorescence imaging was performed using a Cytation-10 system equipped with GEN5 acquisition software. For each experimental group (n = 20-30 worms), photomicrographs were captured at both 10× and 20× magnifications. Image analysis was conducted using ImageJ (NIH, Bethesda, MD), with a consistent region of interest maintained across all conditions to ensure comparability. Quantification of mean fluorescence intensity and subsequent statistical analyses were performed using GraphPad Prism 5.

## Data Analysis

All data were derived from at least three independent experiments, with the number of replicates specified for each endpoint. Results are expressed as mean ± standard error of the mean (SEM). Statistical analyses were performed using ANOVA, and Student’s t-test in GraphPad Prism 5, and significance was defined as *p < 0.05*.

## Acknowledgement

Metabolomics services in support of the research project were provided by the VCU Massey Cancer Center Lipidomics and Metabolomics Shared Resource, supported, in part, with funding from NIH-NCI Cancer Center Support Grant P30 CA016059. S.F.L was supported by NIH R01AG075061 award.

## Supplementary Figures

**Fig. S1.**
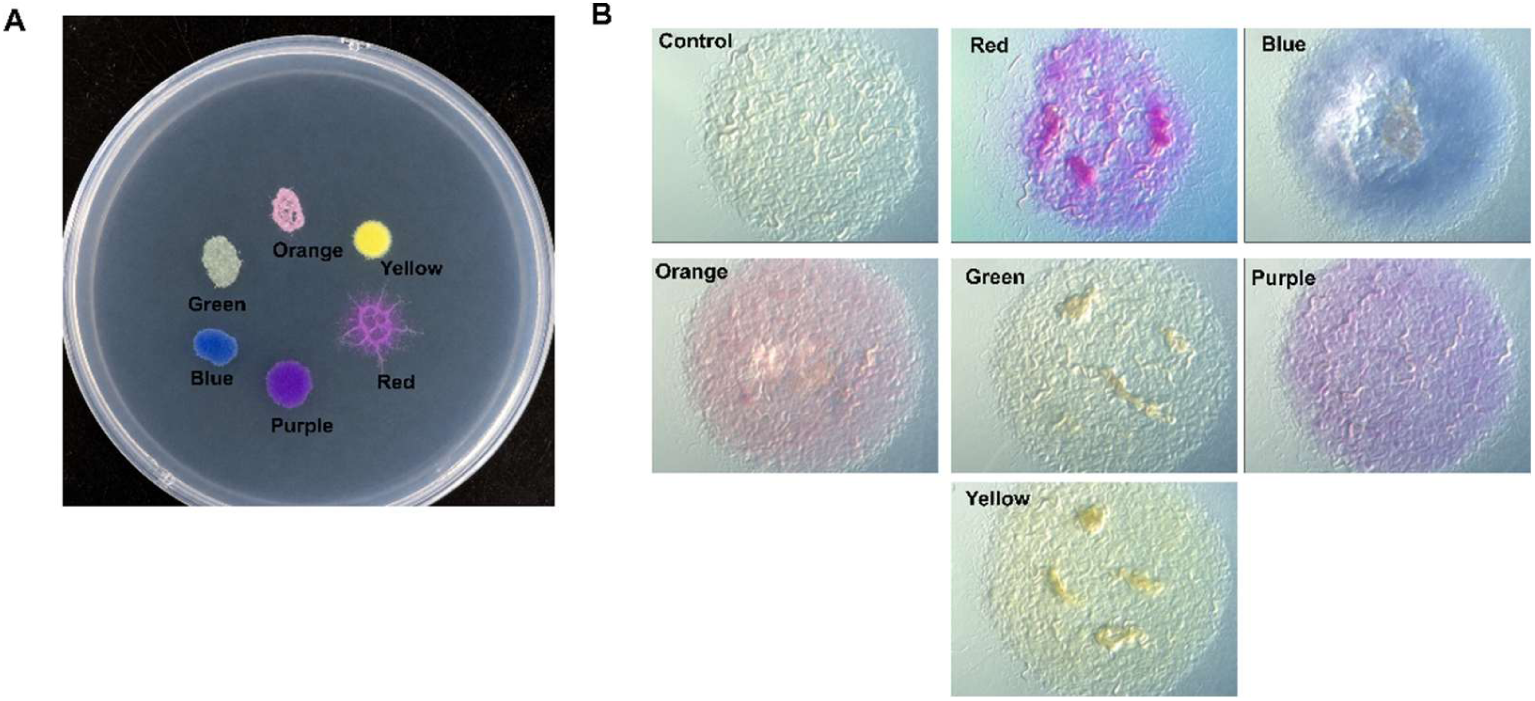
Characterization of bacterial lawn consumption by *C. elegans*. (**A**) Representative image of a bacterial choice assay plate containing spatially separated lawns of bacteria expressing different chromoproteins (red, blue, purple, orange, green, and yellow). Young adult *C. elegans* were placed at the center of the plate and allowed to freely choose among the bacterial lawns. (**B**) Images of individual bacterial lawns following 24 h of feeding by N2 (wild-type) worms. Red chromogenic bacterial lawns exhibited substantially greater depletion than the other chromogenic or control lawns, indicating preferential consumption of red bacteria.

**Fig. S2.**
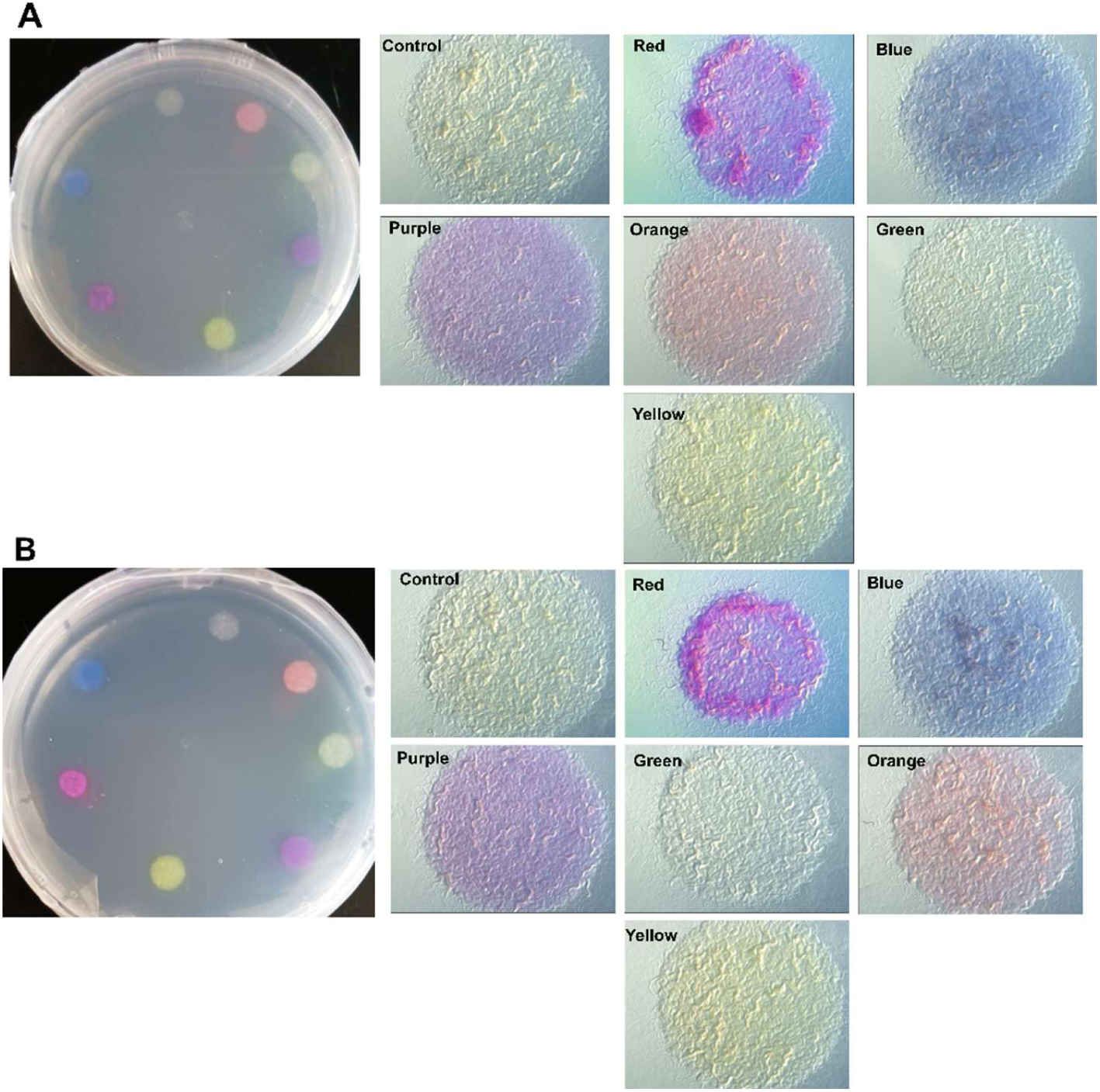
Wild-type worms preferentially consume red chromogenic bacteria independently of ambient light. **(A and B)** Representative images of bacterial consumption by N2 (wild-type) *C. elegans* after 24 h of feeding under ambient light and dark conditions. Assay plates (left) contained spatially separated lawns of control bacteria and bacteria expressing red, blue, purple, orange, green, and yellow chromoproteins. Higher-magnification images (right) show the extent of lawn consumption following feeding. Red chromogenic bacterial lawns exhibited markedly greater depletion than all other lawns under both light and dark conditions, indicating that the preference for red bacteria is maintained in the absence of light.

**Fig. S3.**
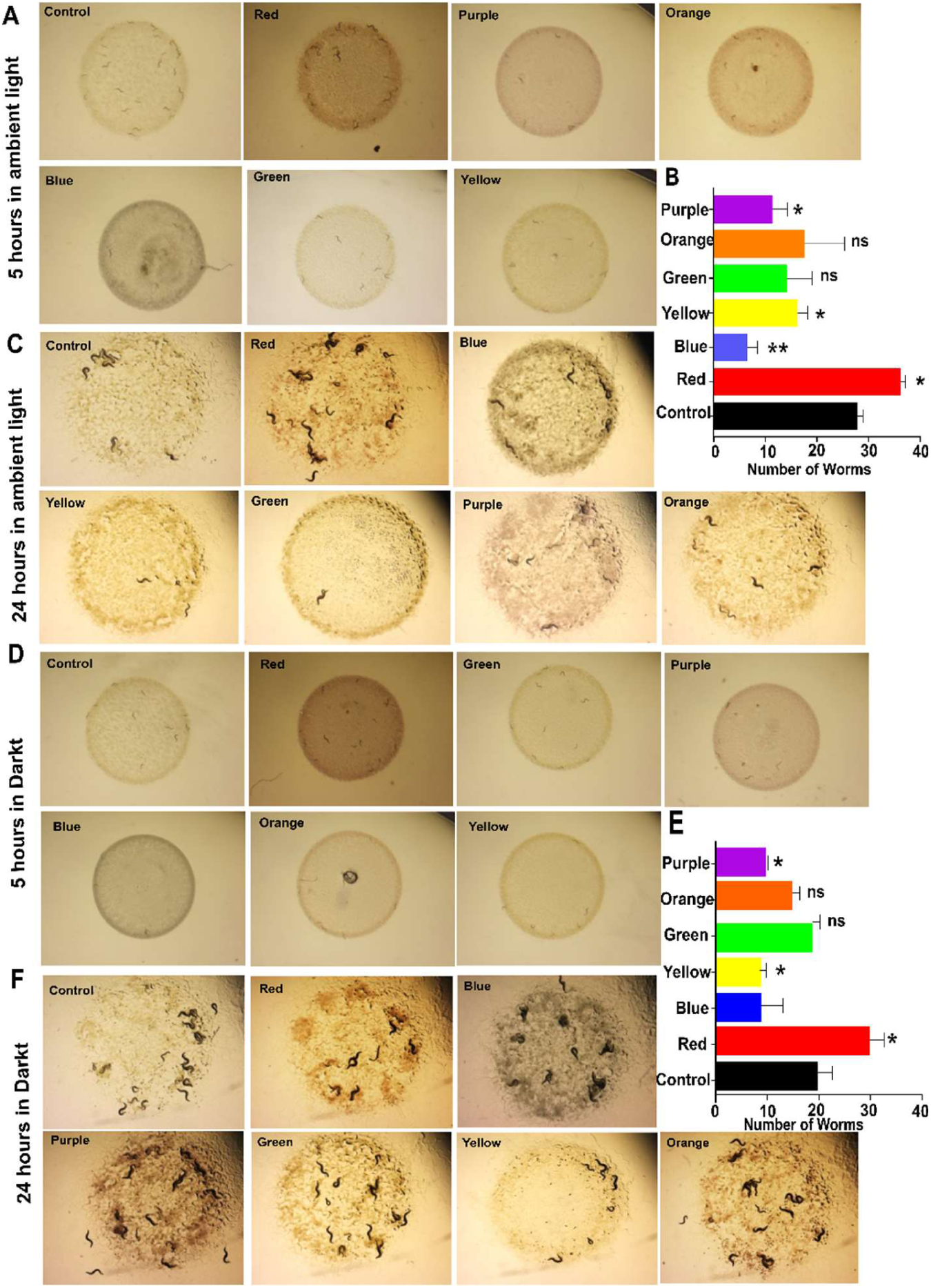
The natural *C. elegans* isolate CX11262 is preferentially attracted to red chromogenic bacteria independent of ambient light. **(A)** Representative images of bacterial choice assays using the natural *C. elegans* isolate CX11262 (Los Angeles, California, USA) after 5 h of incubation under ambient light. **(B)** Quantification of worm distribution on each bacterial lawn under ambient light (*p* = 0.0135 for red; *p* = 0.0052 for blue; *p* = 0.0168 for yellow; *p* = 0.0167 for purple; ns = not significant) student’s t-test. **(C)** Representative images of bacterial choice assays using the natural *C. elegans* isolate CX11262 after 24 h of incubation under ambient light. **(D)** Representative images of bacterial choice assays using the natural *C. elegans* isolate CX11262 after 5 h of incubation in the dark. **(E)** Quantification of worm distribution on each bacterial lawn in the dark after 5 h (*p* = 0.0403 for red; *p* = 0.0380 for yellow; *p* = 0.0370 for purple) student’s t-test. **(F)** Representative images of bacterial choice assays using the natural *C. elegans* isolate CX11262 after 24 h of incubation in the dark. Statistical significance was determined by an unpaired *t*-test compared to the control bacterial lawn.

**Fig. S4.**
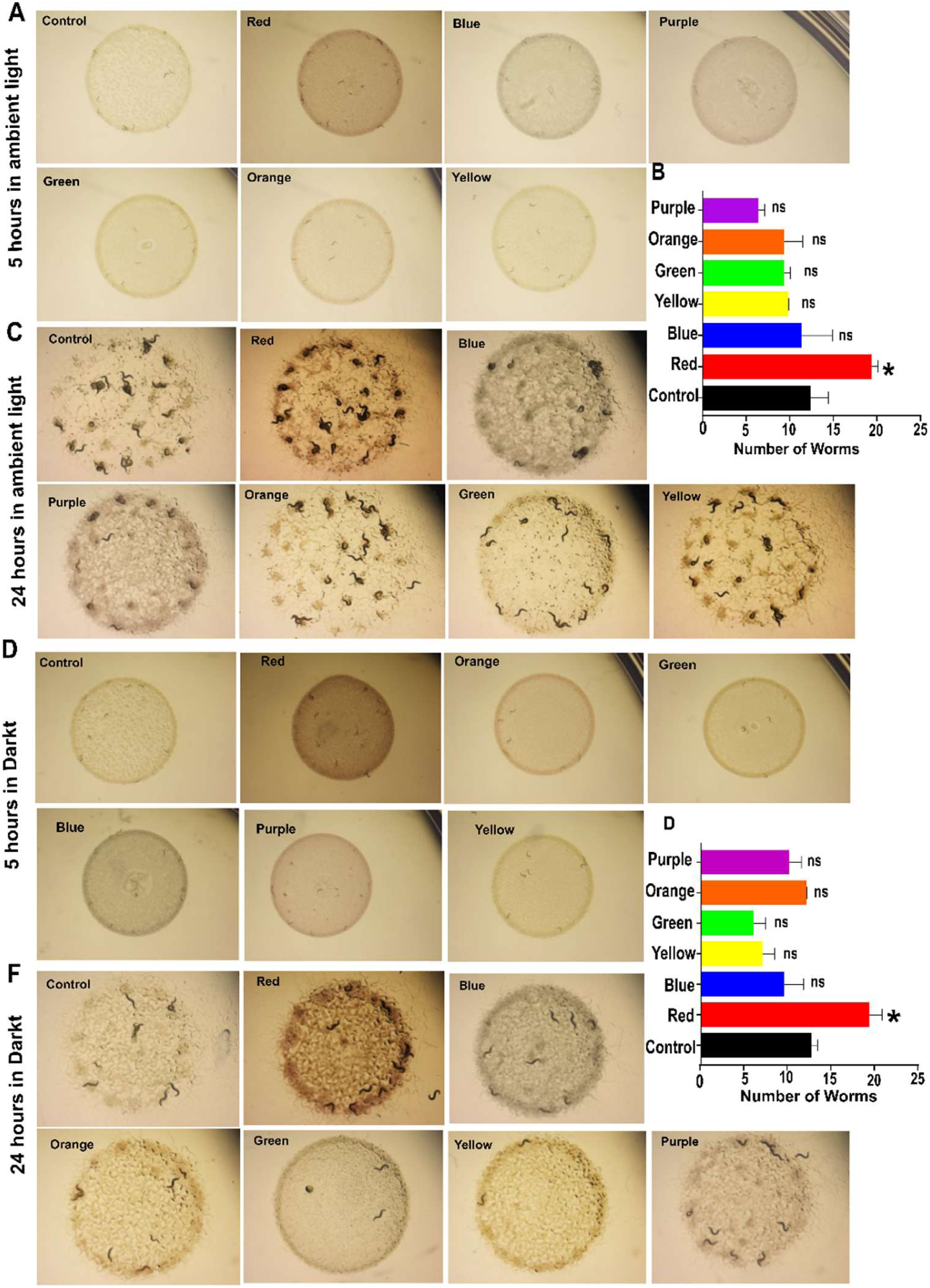
The natural *C. elegans* isolate GXW1 is preferentially attracted to red chromogenic bacteria under both ambient light and dark conditions. **(A)** Representative images of bacterial choice assays using the natural *C. elegans* isolate GXW1 (Wuhan, China) after 5 h of incubation under ambient light. **(B)** Quantification of worm distribution on each bacterial lawn under ambient light (*p* = 0.0124 for red; ns = not significant) student’s t-test. **(C)** Representative images of bacterial choice assays using the natural *C. elegans* isolate GXW1 after 24 h of incubation under ambient light. **(D)** Representative images of bacterial choice assays using the natural *C. elegans* isolate GXW1 after 5 h of incubation in the dark. **(E)** Quantification of worm distribution on each bacterial lawn in the dark after 5 h (*p* = 0.0342 for red) student’s t-test. **(F)** Representative images of bacterial choice assays using the natural *C. elegans* isolate GXW1 after 24 h of incubation in the dark. Statistical significance was determined by an unpaired *t*-test compared to the control bacterial lawn.

**Fig. S5.**
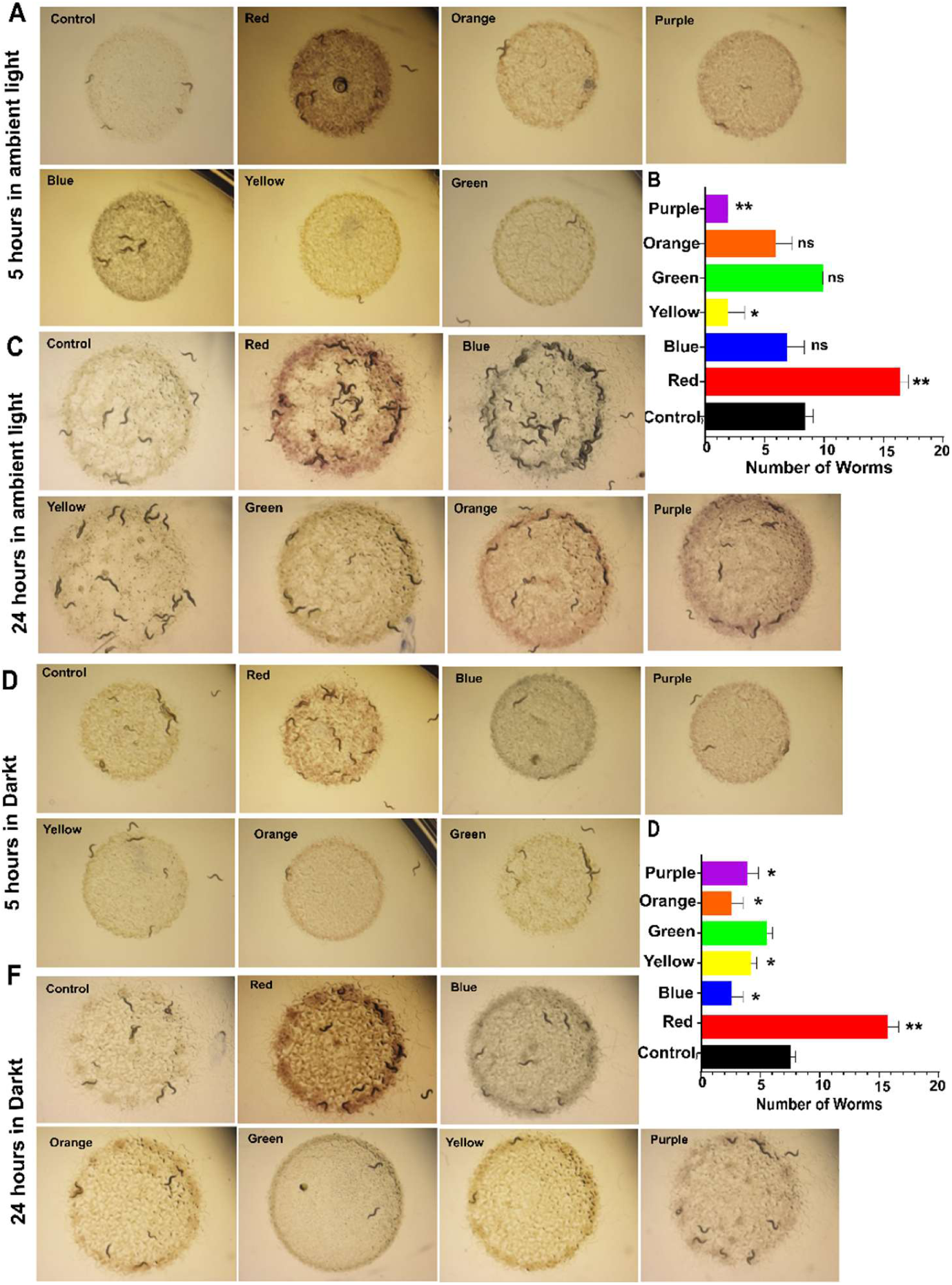
The natural *C. elegans* isolate QX2263 preferentially associates with red chromogenic bacteria under both ambient light and dark conditions. **(A)** Representative images of bacterial choice assays using the natural *C. elegans* isolate QX2263 (Argentina, South America) after 5 h of incubation under ambient light. **(B)** Quantification of worm distribution on each bacterial lawn under ambient light (*p* = 0.0421 for red; *p* = 0.0284 for yellow; *p* = 0.0058 for purple). **(C)** Representative images of bacterial choice assays using the natural *C. elegans* isolate QX2263 after 24 h of incubation under ambient light. **(D)** Representative images of bacterial choice assays using the natural *C. elegans* isolate QX2263 after 5 h of incubation in the dark. **(E)** Quantification of worm distribution on each bacterial lawn in the dark after 5 h (*p* = 0.0076 for red; *p* = 0.0218 for blue; *p* = 0.0194 for yellow; *p* = 0.0218 for orange; *p* = 0.0394 for purple). **(F)** Representative images of bacterial choice assays using the natural *C. elegans* isolate QX2263 after 24 h of incubation in the dark. Statistical significance was determined by an unpaired *t*-test compared to the control bacterial lawn.

**Fig. S6.**
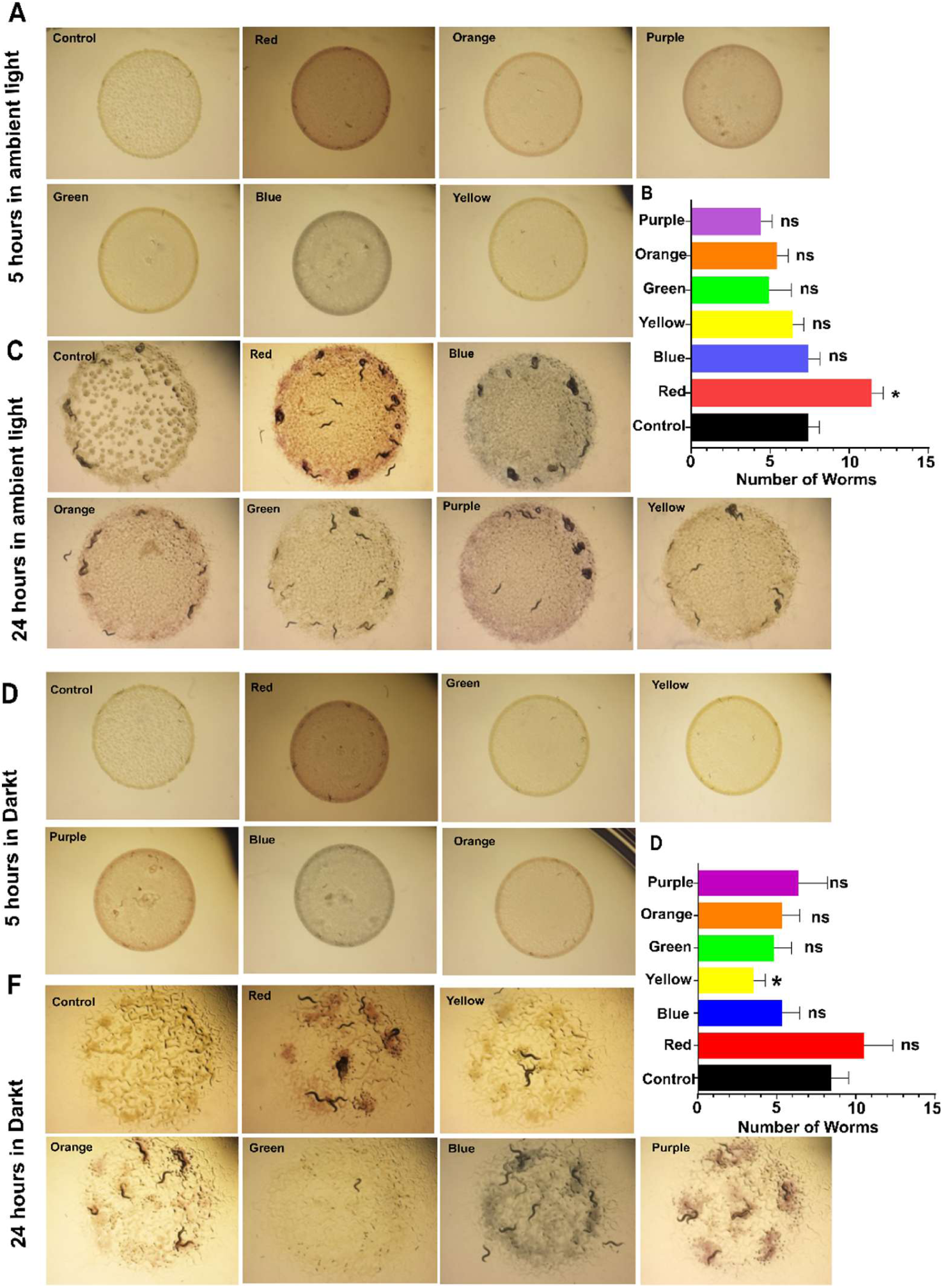
The natural *C. elegans* isolate ED3049 prefers red chromogenic bacteria under both ambient light and dark conditions. **(A)** Representative images of bacterial choice assays using the natural *C. elegans* isolate ED3049 (Ceres, South Africa) after 5 h of incubation under ambient light. **(B)** Quantification of worm distribution on each bacterial lawn under ambient light (*p* = 0.029 for red). **(C)** Representative images of bacterial choice assays using the natural *C. elegans* isolate ED3049 after 24 h of incubation under ambient light. **(D)** Representative images of bacterial choice assays using the natural *C. elegans* isolate ED3049 after 5 h of incubation in the dark. **(E)** Quantification of worm distribution on each bacterial lawn in the dark after 5 h (*p* = 0.041 for yellow). **(F)** Representative images of bacterial choice assays using the natural *C. elegans* isolate ED3049 after 24 h of incubation in the dark. Statistical significance was determined by an unpaired *t*-test compared to the control bacterial lawn.

**Fig. S7.**
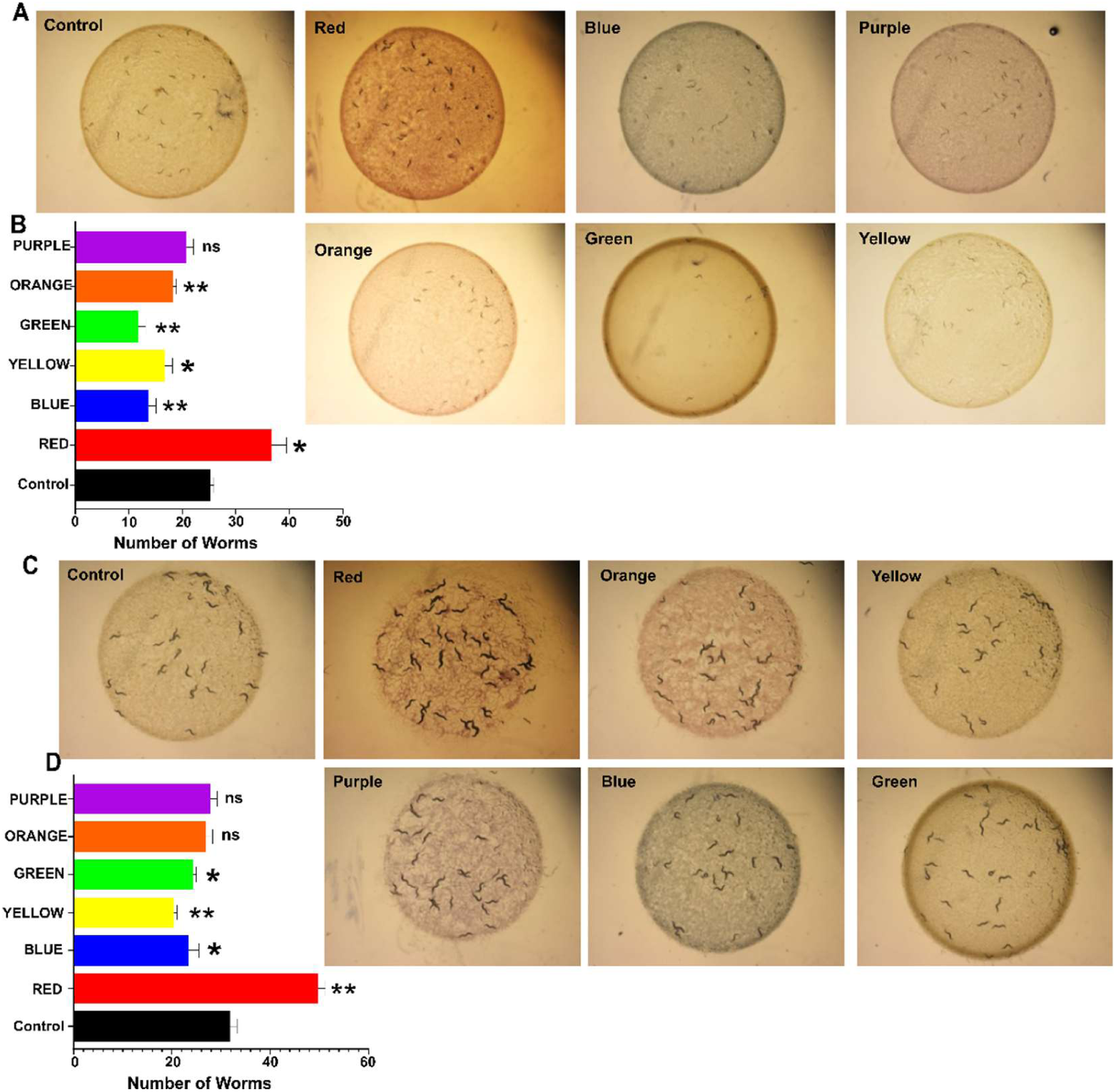
Loss of LITE-1 does not abolish the preference for red chromogenic bacteria. **(A)** Representative images of bacterial choice assays performed with the TQ8245 (*lite-1* null) strain following 5 h of incubation under ambient light. Worms were allowed to choose among a non-chromogenic control and bacterial lawns expressing red, blue, purple, orange, green, or yellow chromoproteins. **(B)** Quantification of worm preference after 5 h under ambient light (*p* = 0.0302 for red; *p* = 0.0092 for blue; *p* = 0.0163 for yellow; *p* = 0.0065 for green; *p* = 0.0095 for orange). Bars represent the mean ± SEM of the number of worms associated with each bacterial lawn. Statistical significance was determined using an unpaired *t*-test relative to the control bacterial lawn. **(C)** Representative images of bacterial choice assays performed with the TQ8245 (*lite-1* null) strain following 24 h of incubation. Worms were exposed to the same set of chromogenic and control bacterial lawns as in (A). **(D)** Quantification of worm preference after 24 h of incubation (*p* = 0.0061 for red; *p* = 0.0419 for blue; *p* = 0.0092 for yellow; *p* = 0.0213 for green). Bars represent the mean ± SEM of the number of worms associated with each bacterial lawn. Statistical significance was determined using an unpaired *t*-test relative to the control bacterial lawn.

**Fig. S8.**
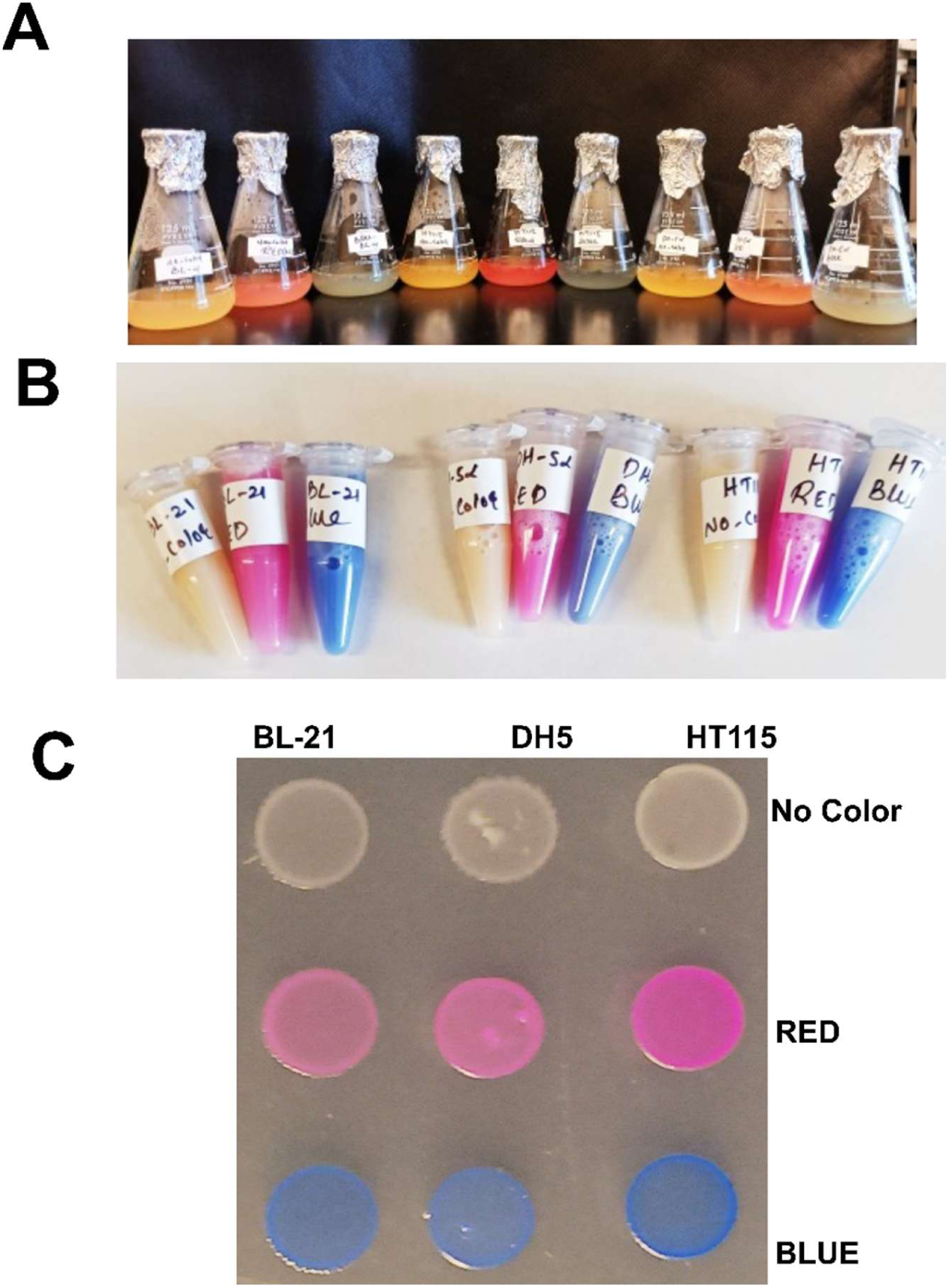
Stable expression of red and blue chromoproteins in multiple bacterial strains used for behavioral assays. **(A)** Representative photographs of liquid cultures expressing chromoproteins in three bacterial strains (BL21, DH5α, and HT115). Bacteria expressing red or blue chromoproteins developed robust pigmentation, whereas control cultures lacking chromoprotein expression remained unpigmented. **(B)** Representative images of bacterial pellets following centrifugation, demonstrating stable accumulation of red and blue chromoproteins in BL21, DH5α, and HT115 cells compared to non-chromogenic controls. **(C)** Representative bacterial lawns of BL21, DH5α, and HT115 expressing no chromoprotein (control), red, or blue chromoproteins after growth on agar plates. Similar pigmentation was observed across all three bacterial backgrounds, confirming consistent chromoprotein expression independent of the bacterial strain.

**Fig. S9.**
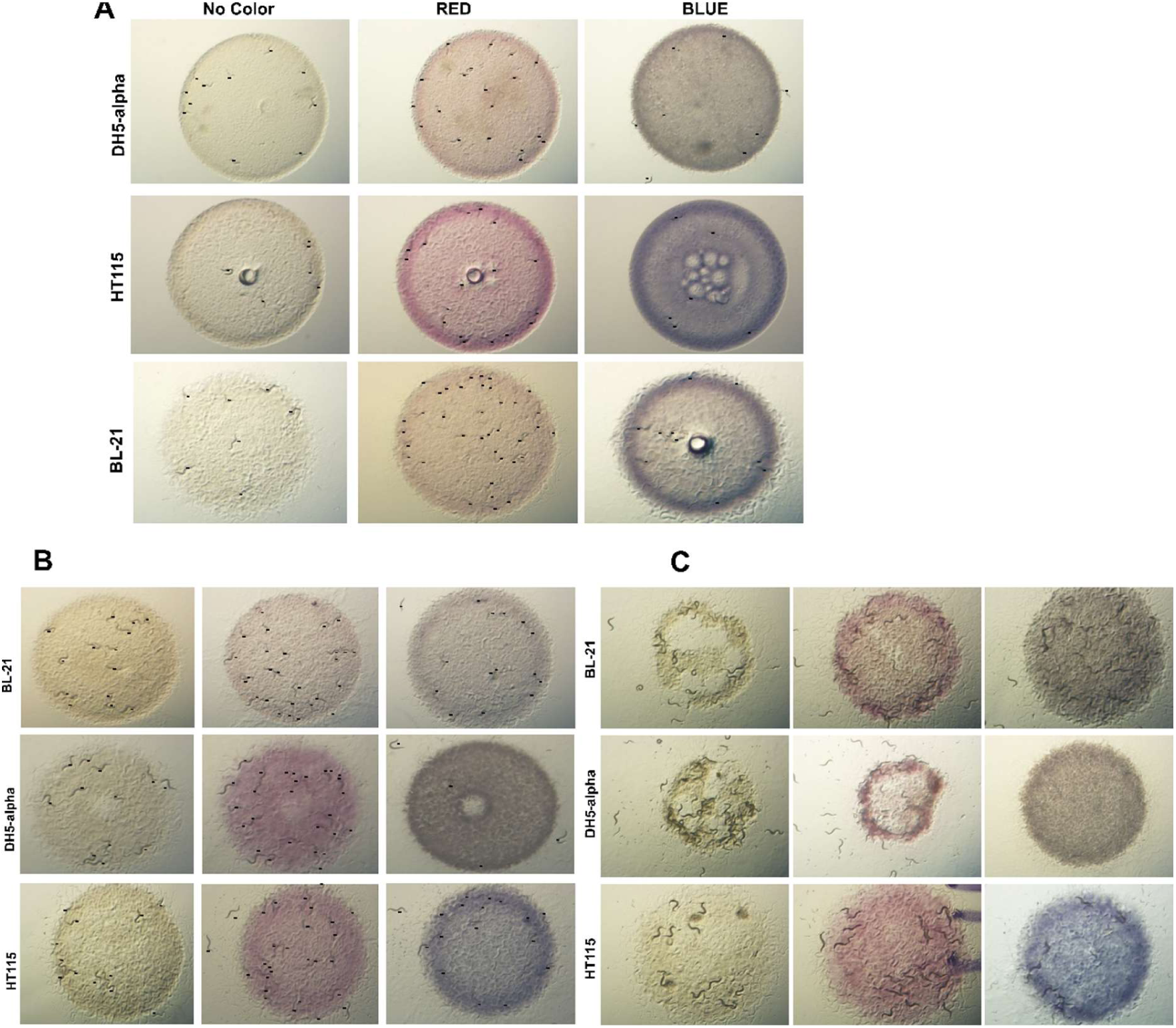
Preference for red chromogenic bacteria is conserved across different bacterial backgrounds and incubation periods. Introductory panels show bacterial attraction assays performed with wild-type (N2) C. elegans using three bacterial backgrounds (DH5α, HT115, and BL21) expressing either red or blue chromoproteins, alongside non-chromogenic controls. **(A)** Representative images of bacterial lawns after 5 h of incubation. Worms preferentially accumulated on red chromogenic bacterial lawns across all three backgrounds, whereas reduced accumulation was observed on blue lawns. **(B)** Representative images after 24 h of incubation. The preference for red chromogenic bacteria was maintained across all bacterial backgrounds following prolonged exposure. **(C)** Representative images after 48 h of incubation. Red chromogenic bacterial lawns exhibited greater consumption than blue or non-chromogenic control lawns across all backgrounds (BL21, DH5α, and HT115), indicating that the preference for red bacteria is independent of the bacterial genetic background and persists throughout extended feeding.

**Fig. S10.**
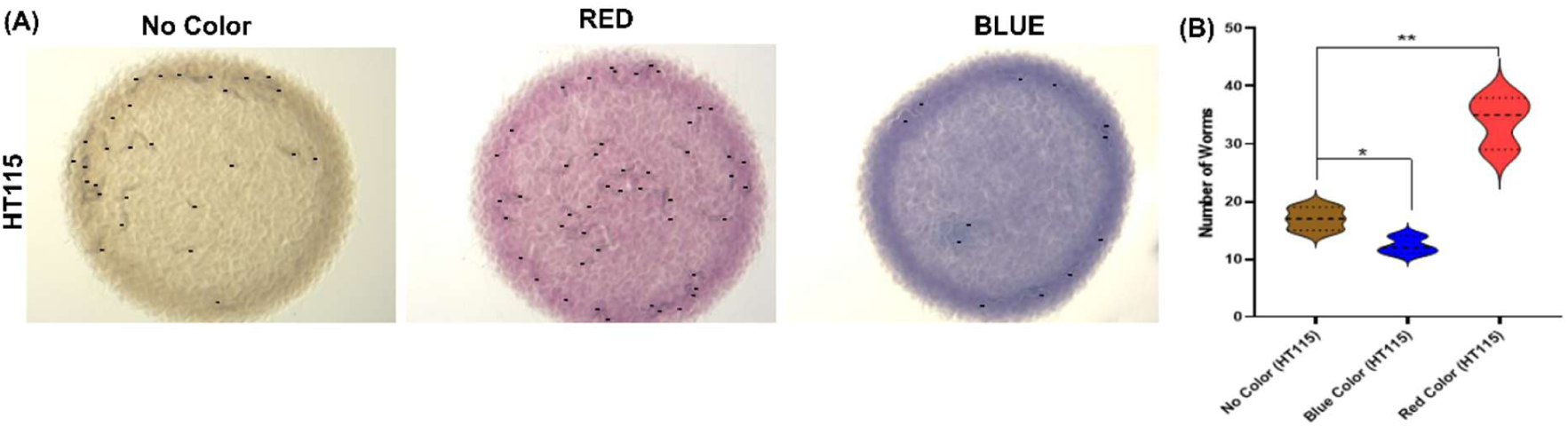
meffRed chromoprotein expressed in HT115 bacteria attracts *C. elegans*. **(A)** Representative images of bacterial attraction assays performed with N2 (wild-type) *C. elegans* using HT115 bacteria expressing the meffRed chromoprotein, blue chromoprotein, or a non-chromogenic control. Worms were allowed to choose for 5 h, after which the number of worms associated with each bacterial lawn was quantified. **(B)** Violin plots showing the distribution of worms associated with each bacterial lawn. The central line indicates the median, and the dashed lines indicate the quartiles. The meffRed-expressing bacterial lawn attracted significantly more worms than either the non-chromogenic control or the blue chromoprotein-expressing lawn. Statistical significance was determined using an unpaired *t*-test (*p* = 0.009 [red]; *p* = 0.043 [blue]).

**Fig. S11.**
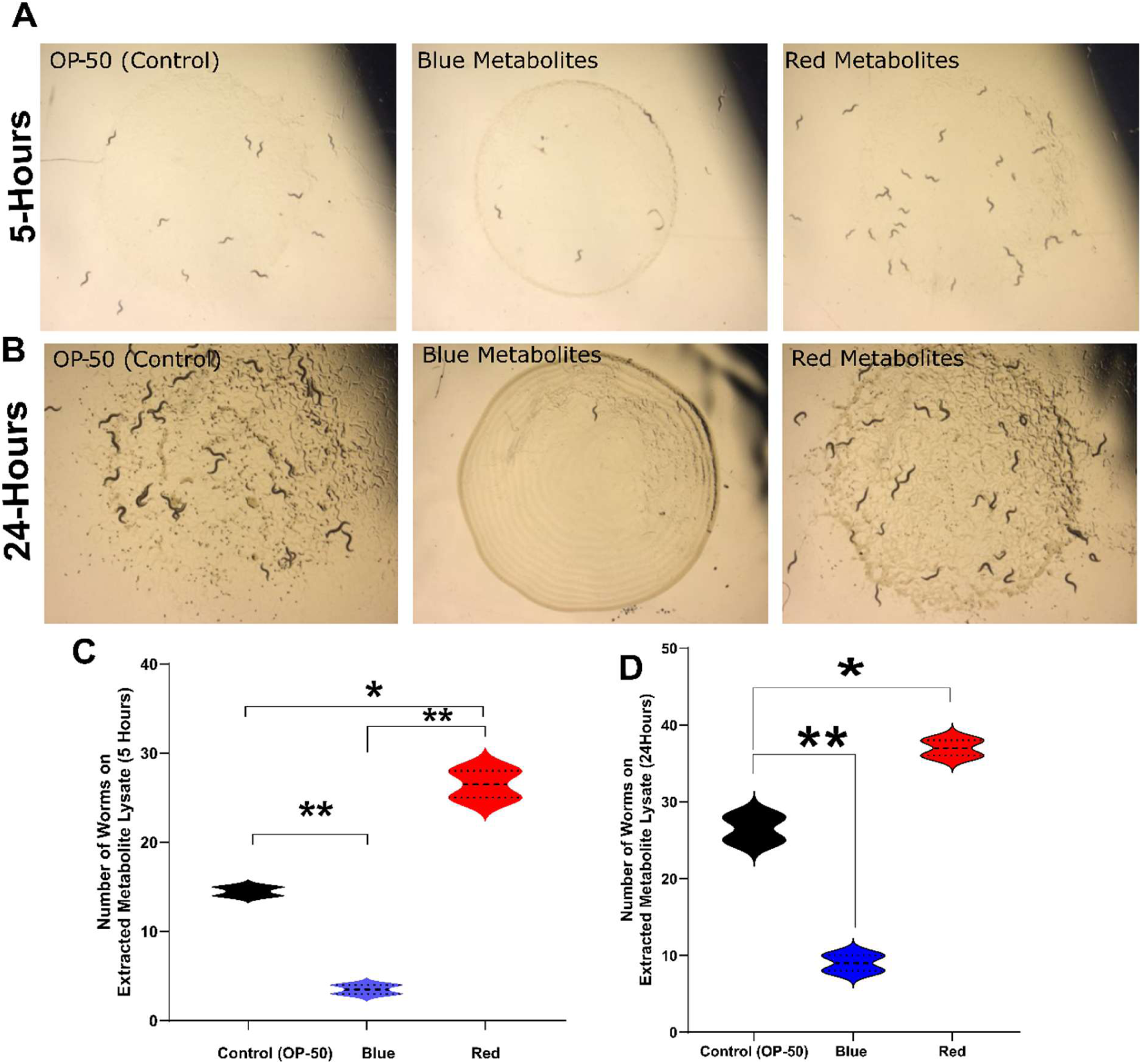
Metabolites extracted from red chromoprotein-producing bacteria are sufficient to promote *C. elegans* attraction. **(A)** Representative images of attraction assays performed with wild-type (N2) worms after 5 h of exposure to *E. coli* OP50 bacterial lawns supplemented with protein-normalized metabolite extracts (1:2 dilution) isolated from red or blue chromoprotein-producing bacteria. OP50 lawns supplemented with extraction buffer alone served as the control. **(B)** Representative images of the same assay after 24 h of incubation. Worms remained preferentially associated with OP50 lawns supplemented with metabolites derived from red chromoprotein-producing bacteria, whereas substantially fewer worms accumulated on lawns containing blue metabolite extracts. **(C and D)** Violin plots showing the distribution of worm numbers associated with each bacterial lawn after 5 h **(C)** (*p* = 0.017 for red; *p* = 0.010 for blue) and 24 h **(D)** (*p* = 0.0027 for red; *p* = 0.0005 for blue). The solid central line indicates the median, and the dashed lines represent the quartiles. Statistical significance was determined using an unpaired *t*-test.

**Fig. S12.**
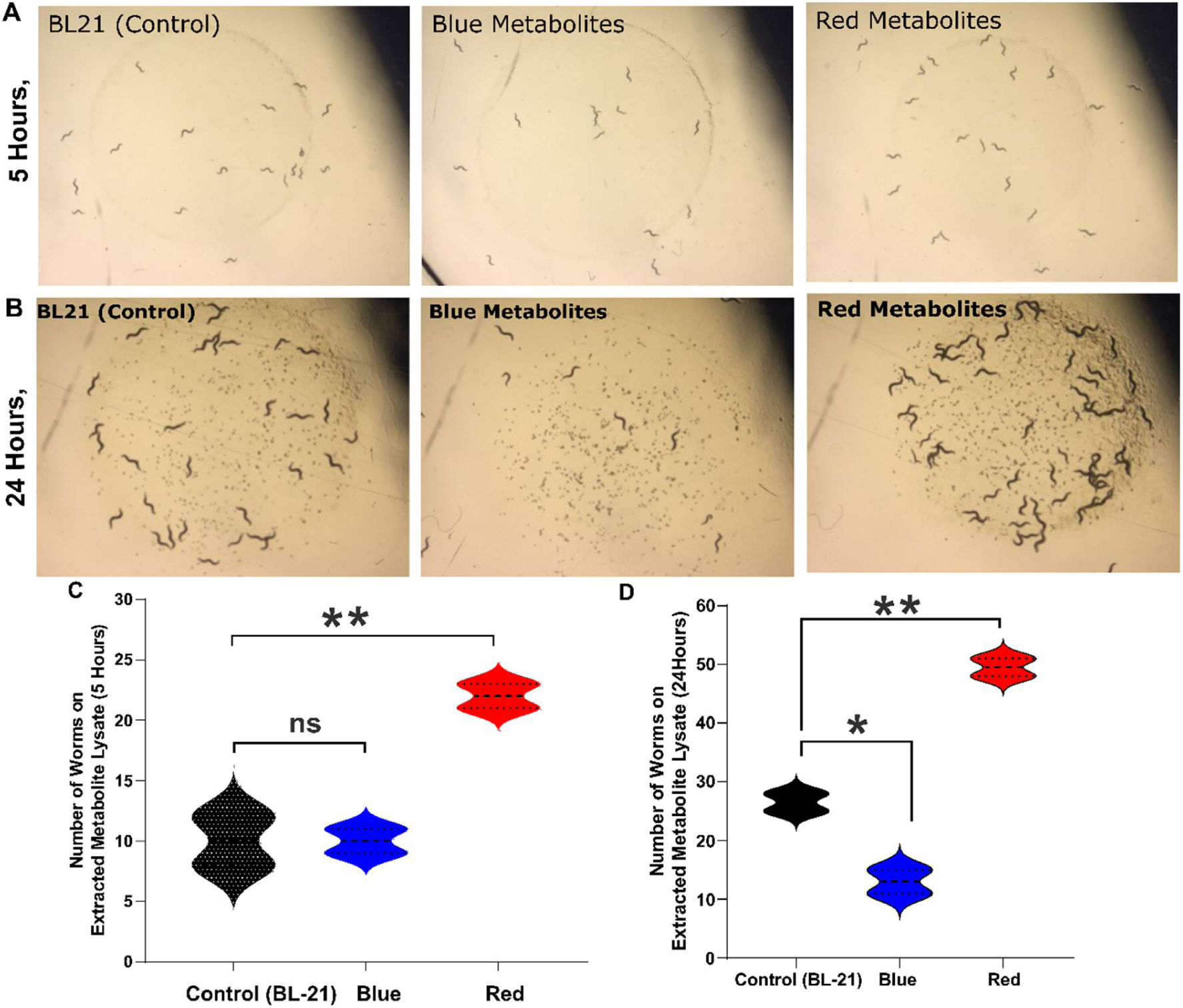
Metabolite extracts from red chromoprotein-producing bacteria enhance *C. elegans* attraction. Representative attraction assays were performed with wild-type (N2) worms using BL21 bacterial lawns supplemented with protein-normalized metabolite extracts (1:2 dilution) isolated from red or blue chromoprotein-producing bacteria. BL21 lawns supplemented with extraction buffer alone served as the control. **(A)** Representative images after 5 h of incubation. Worms preferentially accumulated on BL21 lawns supplemented with metabolites derived from red chromoprotein-producing bacteria, whereas blue metabolite extracts did not significantly alter worm attraction relative to the control. **(B)** Representative images after 24 h of incubation. Red metabolite-treated lawns exhibited greater worm accumulation than either the control or blue metabolite-treated lawns. **(C and D)** Violin plots showing the distribution of worm numbers associated with each bacterial lawn after 5 h **(C)** (*p* = 0.033 for red) and 24 h **(D)** (*p* = 0.0027 for red; *p* = 0.005 for blue). The solid central line indicates the median, and the dashed lines represent the quartiles. Statistical significance was determined using an unpaired *t*-test.

**Fig. S13.**
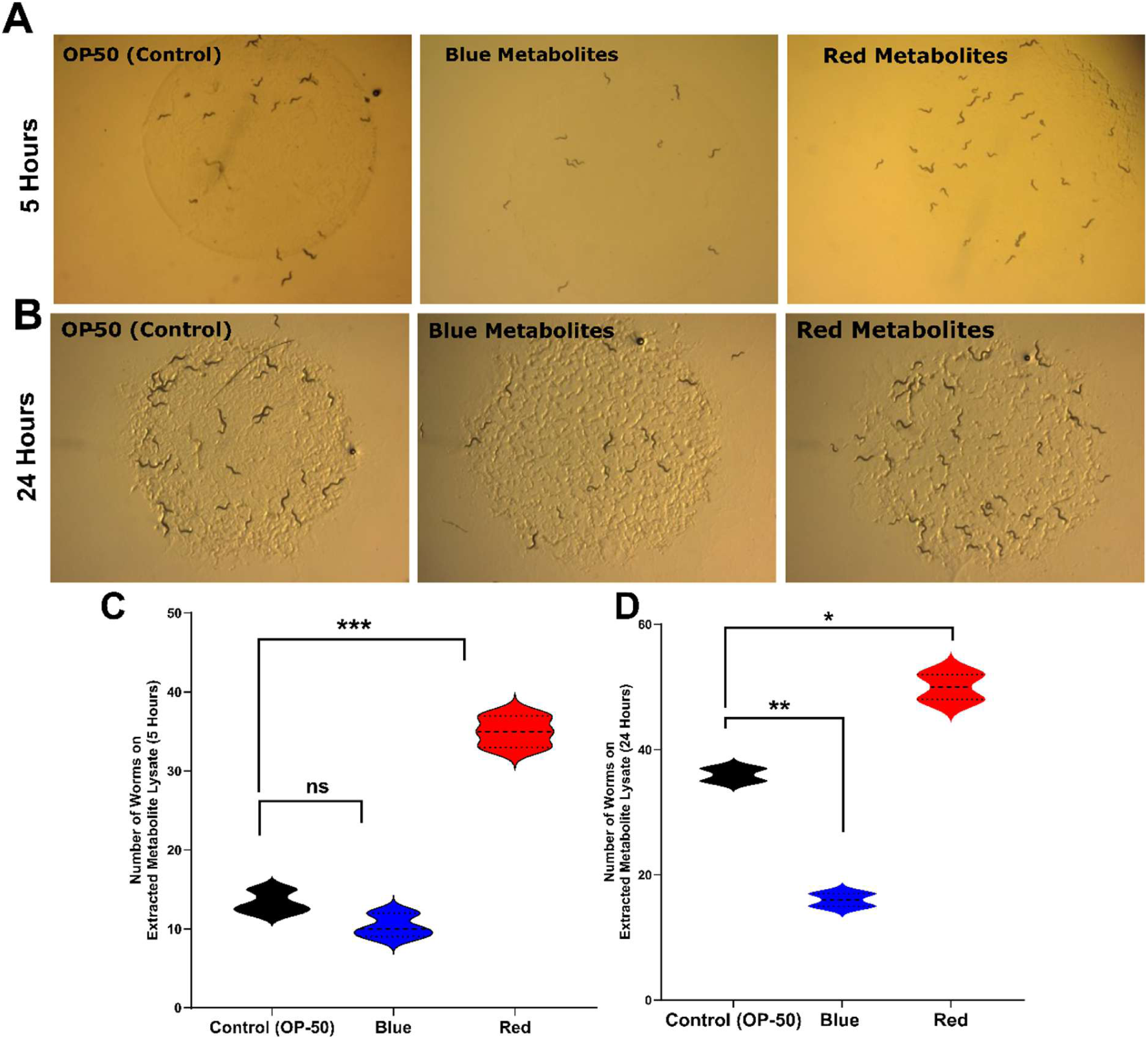
Red chromoprotein-derived metabolites promote attraction of the *lite-1* mutant independent of LITE-1 signaling. Representative attraction assays were performed with the TQ8245 (*lite-1* null) strain using *E. coli* OP50 bacterial lawns supplemented with protein-normalized metabolite extracts (1:2 dilution) isolated from red or blue chromoprotein-producing bacteria (BL21). OP50 lawns supplemented with extraction buffer alone served as the control. (**A**) Representative images after 5 h of incubation. The *lite-1* mutant worms preferentially accumulated on OP50 lawns supplemented with metabolites derived from red chromoprotein-producing bacteria, whereas blue metabolite extracts did not significantly alter worm attraction relative to the control. (**B**) Representative images after 24 h of incubation. Enhanced accumulation of worms on red metabolite-treated lawns persisted after prolonged exposure, while blue metabolite-treated lawns exhibited reduced worm association. (**C and D**) Violin plots showing the distribution of worm numbers associated with each bacterial lawn after 5 h (**C**) (*p* = 0.001 for red) and 24 h (**D**) (*p* = 0.0246 for red; *p* = 0.005 for blue). Statistical significance was determined using an unpaired *t*-test.

**Fig. S14.**
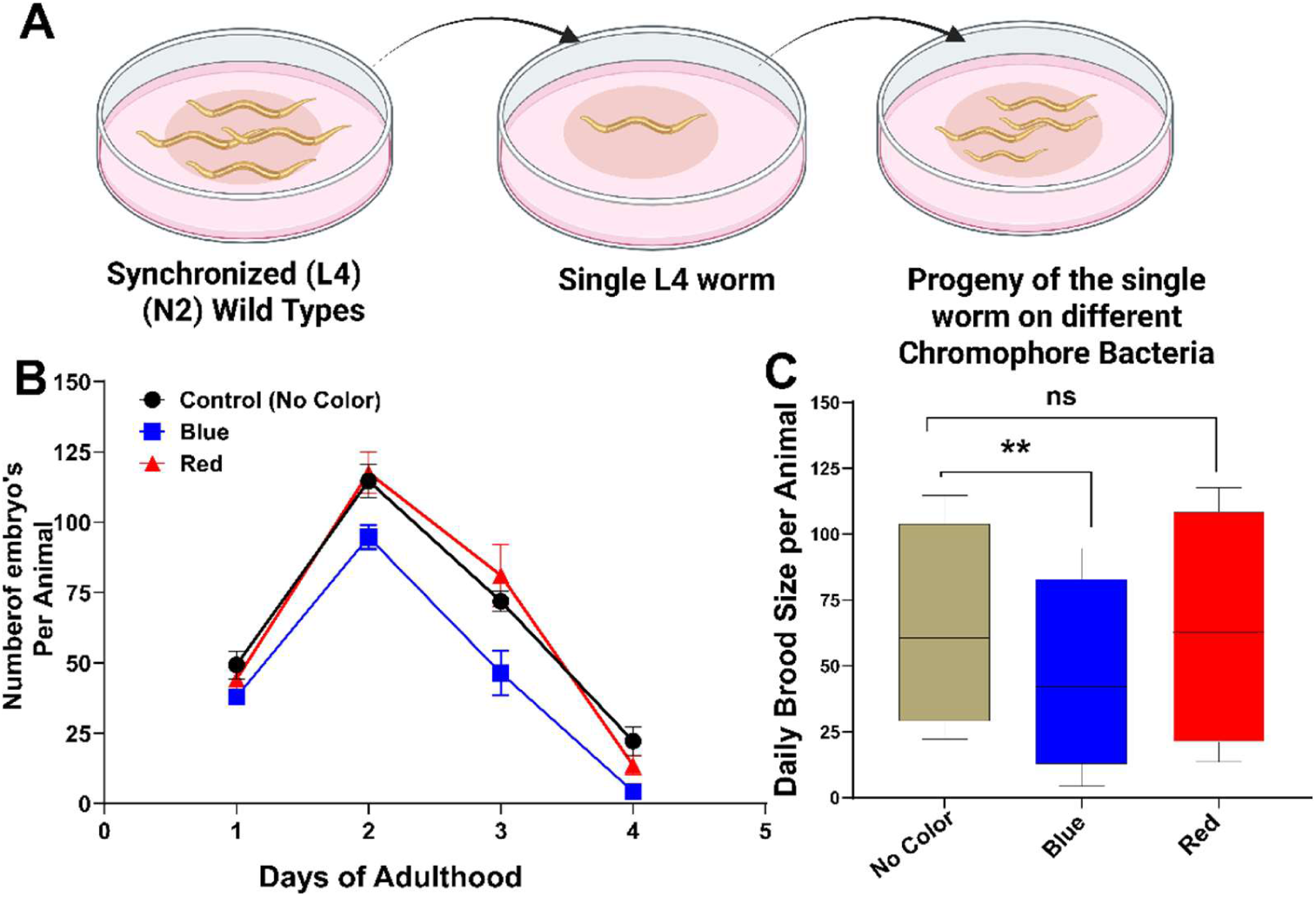
Blue chromoprotein-expressing bacteria reduce fecundity in *C. elegans*. (**A**) Schematic of the experimental procedure used to assess fecundity in N2 (wild-type) worms exposed to non-chromogenic control, red, or blue chromoprotein-expressing HT115 bacteria. Synchronized worms were transferred to the respective bacterial diets at the L4 stage, and reproductive output was monitored daily throughout the reproductive period. (**B**) Daily number of embryos produced per worm on control, red, or blue chromoprotein-expressing bacteria (*n* = 10 worms per condition). (**C**) Quantification of viable progeny produced per worm following exposure to the indicated bacterial diets. Worms exposed to blue chromoprotein-expressing bacteria produced significantly fewer viable progeny than worms fed non-chromogenic control bacteria (*P* = 0.0077; *n* = 10 worms per condition). Data are presented as mean ± SEM. Statistical significance was determined using an unpaired Student’s *t* test.

**Fig. S15.**
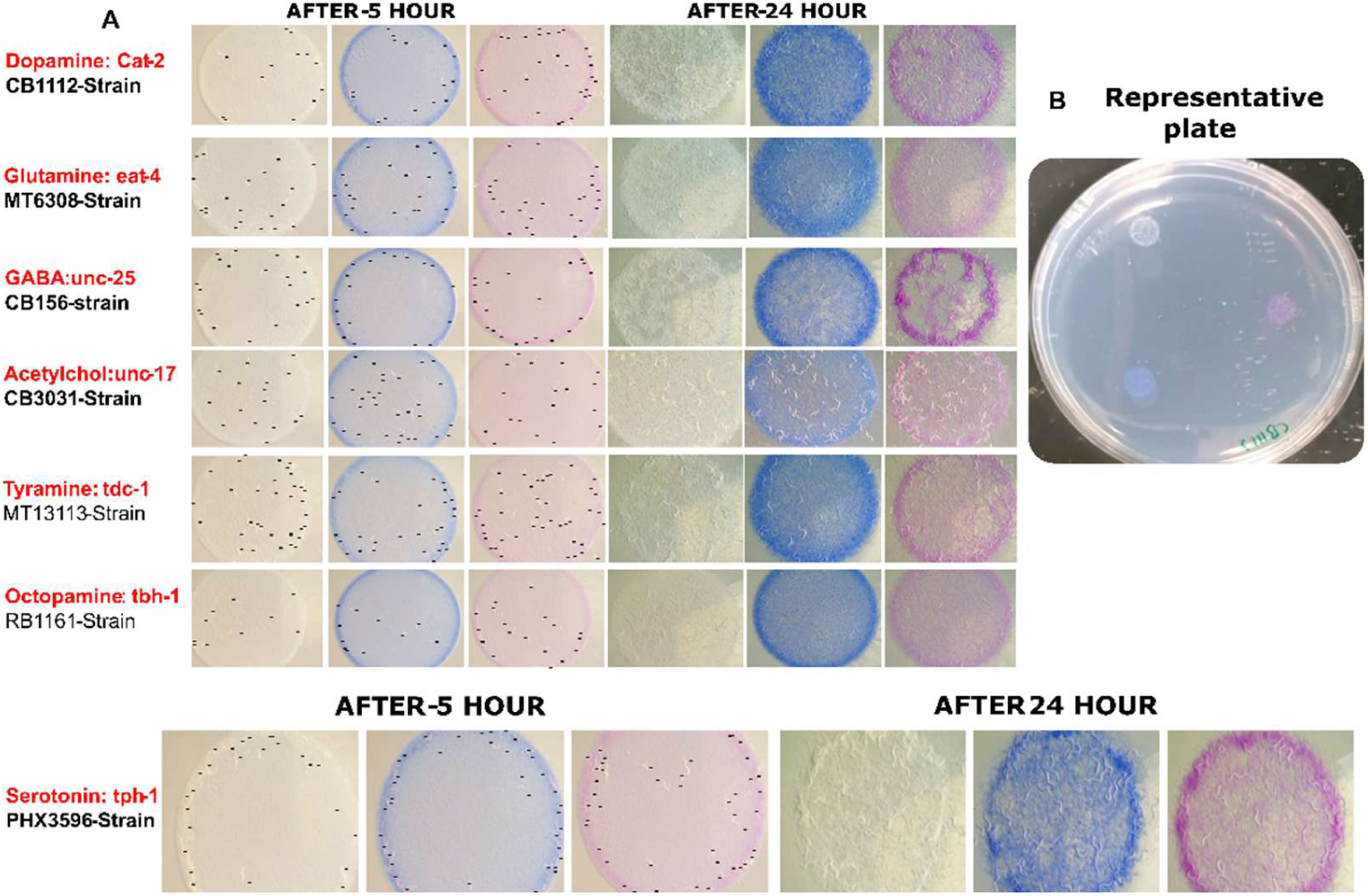
Loss of serotonin signaling identifies a neurocircuit basis for the preference of red chromogenic bacteria. **(A)** Representative images of bacterial choice assays performed with neurotransmitter-deficient *C. elegans* mutants following 5 h and 24 h of incubation. Mutants defective in dopamine (*cat-2*; CB1112), glutamate (*eat-4*; MT6308), GABA (*unc-25*; CB156), acetylcholine (*unc-17*; CB3031), tyramine (*tdc-1*; MT13113), octopamine (*tbh-1*; RB1161), or serotonin (*tph-1*; PHX3596) were allowed to choose among a non-chromogenic control and bacterial lawns expressing red or blue chromoproteins. **(B)** Representative image of a bacterial choice assay plate setup.

**Video S1. Serotonin and neuropeptide signaling are required for red chromophore preference in *C. elegans*.** Representative videos showing the behavioral responses of wild-type N2, serotonin-deficient *tph-1* KO (PHX3596), and neuropeptide-deficient *egl-3* KO (MT150) animals to pure red and blue chromophores. Wild-type N2 worms preferentially migrated toward and accumulated in the red chromophore-containing region, while showing little or no attraction to the blue chromophore. In contrast, *tph-1* KO and *egl-3* KO animals exhibited a marked reduction in red chromophore preference and failed to show the robust accumulation observed in N2 animals. These observations indicate that both serotonergic and neuropeptide signaling contribute to chromophore-specific behavioral preference in *C. elegans*.

