## Supplementary material for "Color-dependent foraging in *C. elegans* integrates chromoprotein photosensitization with bacterial metabolic cues": Table S1

**Table S1. Strain list, used in this study.**

| Strain Name | Genotype | Source |
|---|---|---|
| N2 | Wild type | CGC |
| TQ8245 | lite-1(xu492) X | CGC |
| CB1112 | cat-2(e1112) II) | CGC |
| MT6308 | (eat-4(ky5) III) | CGC |
| RB1161 | tbh-1(ok1196) X. | CGC |
| MT13113 | (tdc-1(n3419). | CGC |
| PHX3596 | (tph-1(mg280) pah-1(syb3596) II). | CGC |
| CB3031 | unc-17(e245) IV; snb-1(e1563) V. | CGC |
| CB156 | unc-25(e156) III. | CGC |
| CB169 | unc-31(e169) IV | CGC |
| MT150 | egl-3(n150) V | CGC |
| GXW1 | WUHAN CHINA ASIA | CGC |
| QX2263 | BUENOS AIRES ARGENTINA SOUTH AMERICA | CGC |
| AB3 | ADELAIDE AUSTRALIA | CGC |
| CB4856 | HAWAII USA NORTH AMERICA | CGC |
| ED3049 | CERES SOUTH AFRICA | CGC |
| RC301 | FREIBURG GERMANY EUROPE | CGC |
| CX11262 | Los Angeles USA AMERICA | CGC |
| PS9050 | flp-4 KNOCKOUT | CGC |
| GR1333 | tph-1::gfp | CGC |
| CZ2475 | juIs145 [flp-13p::GFP + lin-15(+)] II | CGC |
