## Supplementary material for "Color-dependent foraging in *C. elegans* integrates chromoprotein photosensitization with bacterial metabolic cues": Table S2

**Table S2. Chromophores - expressed in *E. coli*.**

| **Chromophore Color** | **Plasmid Names** | **Source/Origin** | **Chromophore Class** |
|---|---|---|---|
| Red | meffRed; eforRed | Montipora efflorescens; Echinopora forskaliana | DsRed-type; Chromoprotein |
| Blue | meffBlue; aeBlue; | Montipora efflorescens; Actinia equina; | Chromoproteins |
| Orange | scOrange | Synthetic | Chromoprotein |
| Purple | tsPurple | Synthetic | Chromoproteins |
| Yellow | fwYellow | Synthetic | Fluorescent (YFP-type) |
| Green | amilGFP; amajLime | Acropora millepora; Anemonia majano | GFP-type |
