## Supplementary material for "Color-dependent foraging in *C. elegans* integrates chromoprotein photosensitization with bacterial metabolic cues": Video-S1

### Slide 1
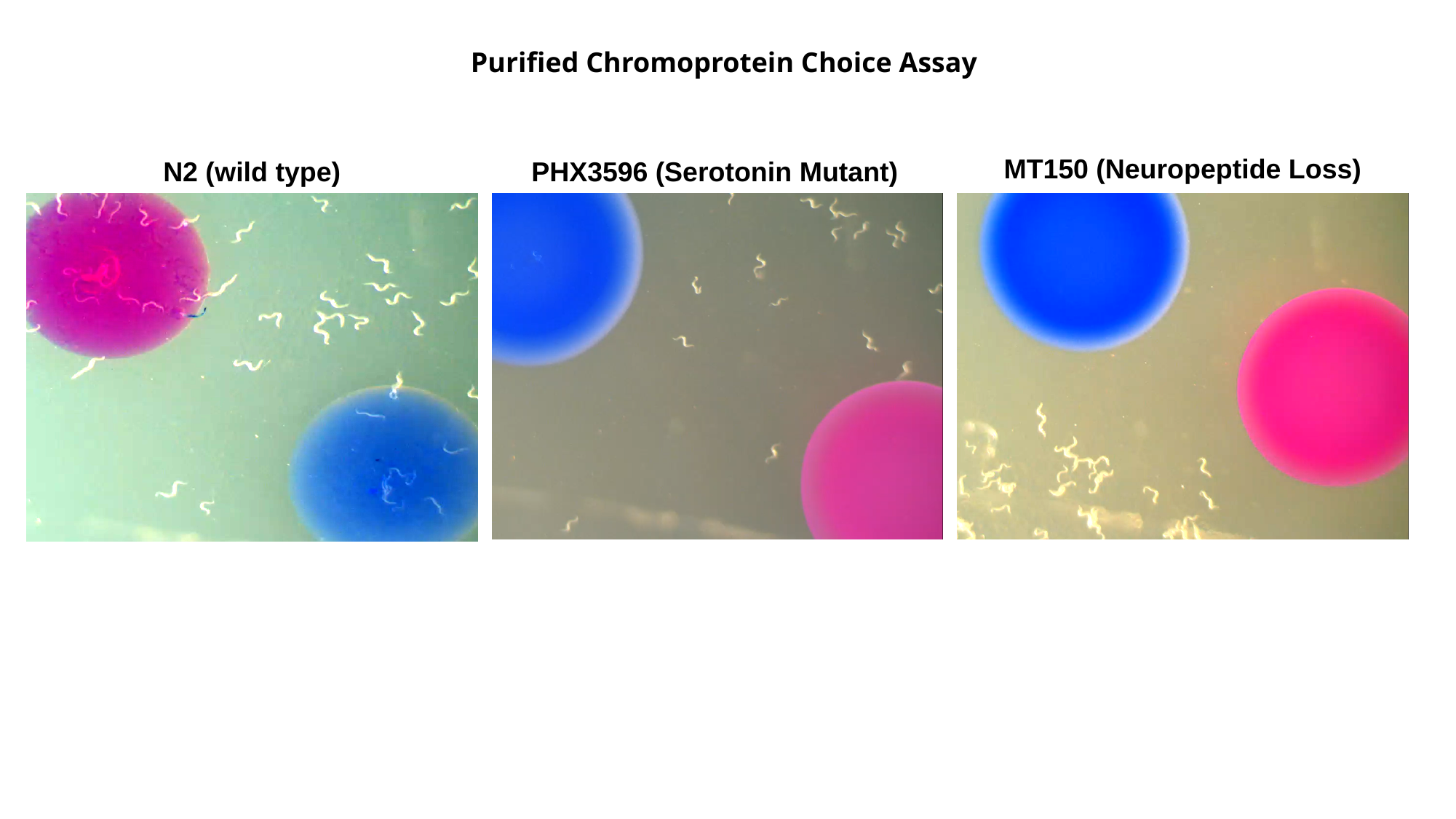

Purified Chromoprotein Choice Assay
MT150 (Neuropeptide Loss)
N2 (wild type)
PHX3596 (Serotonin Mutant)
